# Chromatin architecture and physical constriction cooperate in phenotype switching and cancer cell dissemination

**DOI:** 10.64898/2026.02.05.702638

**Authors:** Pietro Berico, Cody Dunton, Luay Almassalha, Amanda Flores-Yanke, Karla I. Medina, Nicolas Acosta, Tara Muijlwijk, Catherine Do, Soobeom Lee, Sharon N Edmiston, David L. Corcoran, Allison Reiner, Caroline Kostrzewa, Kathy Dorsey, Milad Ibrahim, Shi Qui, Ronglai Shen, Nancy E. Thomas, Amanda W. Lund, Ata S. Moshiri, Iman Osman, Iannis Aifantis, Jane A. Skok, Vadim Backman, Eva Hernando

## Abstract

Phenotypic plasticity is a prominent cancer feature that contributes to metastatic potential and resistance to therapy across multiple cancer types. Cancer cell state transitions have been attributed to transcriptional programs, such as the AP1/TEAD-regulated gene network driving the mesenchymal-like (MES) phenotype. In addition, during dissemination, tumor cells are subjected to variable loads of physical mechanical pressure and constriction across transited tissue, which are thought to impact nuclear molecular crowding. How the interplay between mechanical pressure, global 3D nuclear architecture and transcriptional programs contributes to MES identity and metastatic adaptation remains unclear. Using cutaneous melanoma as a model for early dissemination, we integrate in vitro and in vivo epigenomic profiling with nanoscale imaging of cell lines and patient samples to investigate chromatin organization features underlying the MES phenotype. We find that in MES cells, CTCF is relocated from domain boundaries to regulatory regions of EMT-like genes, leading to reduced insulation, extended topological associated domains (TADs) and increased inter-domain contacts, and de novo formation of chromatin hubs. This conformational rewiring, along with loss of heterochromatin, supports nuclear deformability during invasion and dissemination. Conversely, physical constriction of melanocytic cells induces MES-like chromatin features—including CTCF repositioning and heterochromatin loss— and promotes metastasis in vivo. Similarly, pharmacological inhibition of the heterochromatin mark H3K9me3 triggers MES characteristics and increases invasiveness. These results demonstrate that metastatic competency involves both epigenetic and structural nuclear reprogramming, enabling shifts in gene networks and physical adaptability. Our findings reveal mechanistic links between nuclear architecture and aggressive tumor behavior, identifying potential biomarkers and therapeutic targets to intercept metastatic progression.

## Introduction

Metastasis is the leading cause of death among cancer patients^1–3^. Still, the mechanisms behind cancer dissemination remain largely unknown^4^ and an unmet need to advance therapeutic options. Modern omics technologies have proven that (irreversible) genetic mutations do not fully explain cancer cell metastatic behavior nor patients’ outcome, while ‘reversible’ epigenetic mechanisms can also confer aggressive potential^5, 6^. Cancer cells acquire transient phenotypic states through heritability and plasticity of gene regulatory networks (GRNs) led by different transcription factors (TFs)^7^, as an adaptive response to the tumor microenvironment stimuli^8^. Among the cues, tissue cellular density and matrix composition exert biodynamic forces directly toward tumor cells which can increase molecular crowding within the nuclear compartment^9–11^.

CTCF is the master regulator of nuclear architecture and, together with cohesin complex, nuclear lamina, non-coding RNAs, and other DNA and RNA-binding proteins pack the two meters long human DNA into chromatin, which expands and contracts to regulate gene expression, DNA replication and repair, RNA transcription and splicing^12^. Chromatin is organized into thousands of nanoscale chromatin packing domains (CPDs) that are only vizualied by super-resolution imaging. These are typically up to 200 nm in size and containing 10 kb to 1 Mbp of DNA. These domains represent a distinct organization layer in addition to compartments, TADs and loops that are defined by contacts through chomatin conformation capture or similar approaches, and may function as transcriptional memory elements^13–16^. Not surprisingly, alterations in nuclear architecture can lead to cancer^17^. For instance, genetic and epigenetic aberrations can disrupt compartments, domains and structural loops to rewire cis-regulatory elements (CRE)^18–22^ and establish oncogenic chromatin hubs^23, 24^, composed of CREs simultaneously contacting multiple genes involved in pro-tumorigenic biological processes.

Recently, it has also emerged that chromatin regulation provides nuclear physical structure, mechanical rigidity^25, 26^ and shape^27^ which steer microenvironment adaptability^28, 29^ and may ease tumor cell compression during intra- and extravasation in the metastatic cascade^30^. Concomitantly, mechanical compression can directly rewire 3D chromatin, establishing an intricated bi-directional regulatory feedback between the two processes^28, 31, 32,33^. The amount and maturation of CPDs within the nucleus changes the mechanical stability of a cell; for instance, stable mature domains have been shown to lower the probability of blebbing during cell migration, with predominantly ‘domain-less’ chromatin forming the blebs.^34^ However, the impact of constriction-induced changes on nuclear architecture and phenotype switch during metastasis remains unclear.

A mesenchymal or dedifferentiated cancer cell state (MES), enriched on an epithelial-to-mesenchymal-transition (EMT) program, is thought to be the founder of dissemination in multiple cancer types^35–39^. Skin cutaneous melanoma (SKCM) represents an exquisite model of metastasis because it can disseminate from thin primary lesions^40^ —which already display intra-tumor heterogeneity including MES cells^39, 41, 42^— irrespective of the genetic drivers. In response to microenvironmental cues^43^, melanoma cells ‘switch’ from a differentiated melanocytic-like state (MEL) led by MITF/SOX10 GRNs, to an EMT-like, AP1/TEADs-dependent MES phenotype^44, 45^. However, MITF/SOX10 and AP1/TEADs silencing are insufficient to trigger a complete switch toward a MES state^46, 47^, suggesting other changes in addition to differential gene expression are required ^45, 47–53^. We propose that the bi-directional interplay between nuclear 3D chromatin organization and physical compression critically shapes cancer cell plasticity by impacting both gene expression and mechanical properties. In the context of melanoma, our understanding of global nuclear architecture and its contribution to phenotypic plasticity and metastasis is largely unknown.

Combining multiple omics techniques, we profiled the nuclear physical properties and global 3D chromatin architecture of patient samples, and of a large panel of melanoma cell lines representative of distinct phenotypic states (MEL, intermediate —INT— and MES) or MEL upon silencing of MITF and SOX10 (siMS), the key GRNs of the MEL state. Compared to MEL, siMS and INT cells, we observed a dramatic rewiring of 3D chromatin in MES cells and a preferential binding of CTCF to CRE of EMT-related genes and chromatin hubs over TAD boundaries. Accordingly, silencing of CTCF re-activates MEL-like GRNs and disrupts the activity of a subset of chromatin hubs, some of which support MES cell migration. While melanoma states exhibit extensive differences in nuclear topology, these only result in limited transcriptional changes, suggesting that MES-like 3D chromatin features confer additional biological properties contributing to metastasis beyond gene expression, such as nuclear deformability and flexibility. Supporting this hypothesis, we find MES cells display higher nuclear flexibility and lower stiffness by mathematical polymer simulations, Partial Wave Spectroscopy (PWS) reduced H3K9me3 packing scale domains by spectroscopic single-molecule localization microscopy (sSMLM) and increased cis-long interactions by HiC. Strikingly, these molecular features are partially adopted by MEL cells undergoing constriction during transwell migration or upon H3K9me3 inhibition, which are both independently able to increase their metastatic potential. Altogether, our study shows that nuclear architecture and cell compression co-regulate gene expression and mechanical properties unleashing the MES identity and metastatic capacity.

## Methods

### Cell culture

All melanoma cell lines were grown at 37°C in 5% CO_2_ and routinely tested for mycoplasma contamination using MycoAlert® PLUS Mycoplasma Detection Kit (Lonza, #LT07-703). SKMEL-5, SKMEL-28, SKMEL-239 and SKMEL-147 were kindly provided by Alan Houghton (Memorial Sloan-Kettering Cancer Center, New York, NY, USA); 501mel were purchased from Yale University, MM cell lines (MM057, MM074, MM087, MM029 and MM099) were obtained from Prof. Ghanem Ghanem (Institute Jules Bordet, Bruxelles, BE). MM047 cells were obtained from Frédéric Coin and Irwin Davidson (IGBMC, Illkirch, France). 501mel and SK-MEL cell lines were grown in 10% Fetal Bovine Serum (FBS) in DMEM and penicillin-streptomycin (PS). MM cell lines were grown in Ham-F10 supplemented with 10% FBS, PS and 2mM L-Glutamine.

Cells were transfected with 25nM of siNTC (Horizon Discovery, #D-001810-10), siMITF (Horizon Discovery, #L-008674-00) and siCTCF (Horizon Discovery, #L-020165-00) using Lipofectamine RNAiMAX (Thermo Scientific Scientific, 13778150) following manufacturer’s instructions.

### Transwell migration assay

For migration under constriction, 1x10^6^ 501mel and MM099 were seeded into cell culture inserts with 3.0μm pore size (Corning, #353091) into a dedicated 6 well plate (Corning, #353502) and grown in complete media. 72hrs later, cells were gently detached from both sides of the insert using trypsin and processed for HiC, CUT&Tag (CTCF, H3K9me3 and H3K27ac), ATAC and RNAseq.

To assess the effects of H3K9me3 perturbation, 1-2x10^5^ 501mel and SKMEL-28 were seeded into cell culture inserts with 8.0μm pore size (Corning, #353097) placed into a dedicated 24 well plate (Corning, #353504) and grown in complete media supplemented with DMSO or 5nM Chaetocin (MedChem Express, #HY-N2019) for 12-24hrs. Next, inserts were rinsed twice in PBS and fixed with 4% PFA-PBS solution (Electron Microscopy Sciences, #15733-10) for 10min at room temperature. Next, inserts were stained in 0.2% crystal violet solution for 15min at room temperature, rinsed in distilled water by simultaneously cleaning the inner side with a cotton swab. After drying at room temperature, inserts were imaged using a digital brightfield microscope (Invitrogen, EVOS core) and quantified with ImageJ.

### ChIP and HiC

Melanoma cell lines were double crosslinked for 30min in 25nM EGS (Thermo Scientific Scientific, #21565) and for 10min in 1% FA (Sigma-Aldrich, #47608) at room temperature and then processed for ChIP and HiC.

For ChIP, 1x10^7^ cells were lysed in ChIP lysis buffer (CLB = 10mM Tris HCl pH8, 100mM NaCl, 1mM EDTA pH8, 0.5mM EGTA pH8, 0.1% Sodium Deoxycholate, 0.5% N-lauroylsarcosine, 1% Triton X-100) and sonicated with Diagenode Bioruptor for 15-30 cycles (30” ON +30” OFF max intensity). 5ug/IP of H3K27ac antibody (Active Motif, #91193), H3K9me3 (Active Motif, #61013) or IgG control (Cell Signaling Technology, #3900) were prebound to 50ul of Protein A/G magnetic beads (Pierce Thermo Scientific Scientific, #88803) in PBS+BSA 0.5% for 1hr @RT. Chromatin was diluted to 1ml in CLB and magnetic beads-antibody complex was added to each IP ON at 4°C on revolver. IP were washed twice in Wash Buffer I (WBI, 20mM Tris HCl pH7.5, 150mM NaCl, 2mM EDTA pH8, 0.1% SDS, 1% Triton X100), Wash Buffer II (WBII, 10mM Tris HCl pH8, 250mM LiCl, 1mM EDTA, 1% Triton X100, 0.7% Sodium Deoxycholate) and TET Buffer (10mM Trist HCl pH8, 1mM EDTA, 0.2% Tween20) and eluted in Elution buffer (EB, 10mM Tris HCl pH8, 300mM NaCl, 5mM EDTA pH8, 0.5% SDS, 10ug RNAse A, 100ug proteinase K) 30min at 37°C + ON at 55°C. IP elutions were purified using MinElute Reaction Cleanup kit (Qiagen, #28204) and libraries were prepared using NEBNext Ultra II DNA Library Prep Kit for Illumina (NEB, #E7600S) following manufacturer’s instructions.

For HiC, 2.5x10^6^ cells were processed using Arima-HiC+ kit (Arima Genomics, #A101020) following the manufacturer instructions. HiC libraries were generated using Arima-HiC+ kit (Arima Genomics, #A510008, #A303011) following manufacturer’s instructions.

ChIP and HiC libraries were pooled together at 2nM concentration and sequenced paired-end 2x150bp using Illumina NovaSeqX+.

### HiC in DNA from FFPE tissues

Between 5-10 Formalin Fixed Paraffin Embedded (FFPE) patient tumor tissue sections of 10μm per sample were cut and pooled in a 1.5mL tube for de-waxing and re-hydration using xylene, ethanol and molecular grade water. Next, FFPE samples were processed with the Arima-HiC+ FFPE Kit (Arima Genomics, #A101060) following manufacturer’s instructions. HiC FFPE libraries were generated using Arima-HiC+ kit (Arima Genomics, #A510008, #A303011) following manufacturer’s instructions.

### Omni-ATACseq

5x10^5^ cells were harvested and assessed to not have more than 15% of cell death using trypan blue. Next, dead cells were additionally removed treating cells with 200U/ml of DNAse (Worthington, #LS002007). Cells were resuspended in 500ul of cold buffer 1 (10mM Tris-HCl pH 7.5, 10mM NaCl, 3mM MgCl_2_, 0.1% NP-40, 0.1% Tween-20) and the equivalent volume for 5x10^4^ cells (50ul) was kept for the next steps. A final concentration of 0.1% of digitonin was added to the cells and incubated on ice for 3 minutes. Digitonin permeabilization was quenched adding 1ml cold buffer 2 (10mM Tris-HCl pH 7.5, 10mM NaCl, 3mM MgCl_2_, 0.1% Tween-20). Afterward, cells were pelleted and resuspended in 50ul of transposition mixture (20mM Tris HCl pH 7.5, 10mM MgCl_2_, 20% dimethyl formamide, PBS, 1% digitonin, 10% tween-20) supplemented with 2.5-5ul of Transposase (Illumina, #20034197) and incubated in thermomixer at 37°C for 30 minutes with 1000rpm. Transposition mixture was purified using DNA Clean and Concentrator-5 kit (Zymo, #D4014) for library preparation. 20ul of transposed DNA was processed for Nextera library using the same protocol described in Schmidl work^54^. OmniATAC libraries were pooled together at 2nM concentration and sequenced paired end 2x150 using Illumina NovaSeqX+.

### CUT&Tag

For CTCF genomic occupancy was assessed using CUT&Tag-IT^TM^ (Active Motif, #53160) following manufacture instructions. Briefly, 1.5-5x10^5^ cells were captured with concanavalin A beads and 1ug of IgG control (Cell Signaling Technology, #3900) or CTCF (Active Motif, #61311) antibodies were incubated 2hrs at 4°C on revolver in cell lysis buffer. For H3K9me3 and H3K27ac Pre and Post migration, 1ug of antibodies H3K9me3 (Active Motif, #39062) and H3K27ac (Active Motif, #39133) were incubated overnight at 4°C on revolver in cell lysis buffer. Afterward, 1ul of guinea pig anti-rabbit antibody was added to the reaction followed by 1hr of tagmentation at room temperature with 1ul of Tn5 transposase enzyme. Tagmentation was stopped with EDTA, SDS and proteinase K incubation for 1hr at 55°C. Tagmented DNA was purified using DNA purification reagents within the kit. Finally, tagmented DNA was processed for Nextera library preparation using i7 and i5 indexed primers and Q5 TAQ polymerase. CUT&Tag libraries were pooled together at 2nM concentration and sequenced paired end 2x50 using Illumina NovaSeqX+.

### RNAseq

Total RNA libraries were prepared using automated TruSeq stranded total RNA and RiboZero Gold kit (Illumina, #RS-122-2301) according to the manufacturer’s instructions. RNA libraries were sequenced paired end 2x50 using Illumina NovaSeqX+.

### Methylation array

DNA methylation profiling was performed using the genome-wide Infinium MethylationEPIC array (Illumina) as described previously^55^. Briefly, DNA (250ng/ sample) from melanoma cell lines was extracted with QIAamp DNA Micro Kit (Qiagen, #56304) and bisulfite-treated using EZ DNA Methylation GOLD kits (Zymo Research, #11-335) according to manufacturer’s protocol. BeadArray processing and scanning was performed in the UNC-CH Mammalian Genotyping Core. Methylated and non-methylated control samples (Zymo Research, #50-444-307) were included in the experiment. BeadArray data was assembled using Illumina GenomeStudio methylation module (v3.2). Background intensity from negative controls was subtracted from each data point.

### Tumor dissociation

Freshly dissected tumors from mice were dissociated at single cell by first chopping the tissue in a petri dish with a scalpel. Then, whole tumors were resuspended in 4mL/gr of HBSS (Sigma-Aldrich, #H6648) supplemented with 750U/ml of Collagenase I (Sigma-Aldrich, #C2674) and 625U/ml of DNAse I (Worthington, #LS002138) and incubated for 50min at 37°C upon strong shaking. Cell suspension was sequentially filtered through a first 70μm cell strainer (Fisher Scientific, #22363548) and then a 40μm cell strainer (Fisher Scientific, #22363547). Next, single cell suspensions were depleted of immune and dead cells by flow cytometry sorting (Sony Biotechnology, SY3200 HAPS) using PE-conjugated CD45 (BioLegend, mouse #111103) and DAPI staining.

### PIPseq

Before processing, cell number and viability were assessed using a TC20 Cell Automated Cell Counter (BioRad). Briefly, cells were partitioned into PIPs (PIPseq T20 3’ Single Cell RNA Kits, V4.0PLUS, Fluent BioScience) with subsequent cDNA generation and final library preparation following manufacturer’s instructions. Sample input varied from ∼80,000 to 200,000 per reaction. cDNA quality and concentration were evaluated on a BioAnalyzer 2100 using a High Sensitivity DNA Kit (Agilent Technologies). Final gene expression libraries were amplified between 6 and 15 cycles and visualized on an Agilent TapeStation 4200 using High Sensitivity D1000 ScreenTape (Agilent Technologies). Sequencing was performed using a NovaSeq X+ 10B 200 Cycle Flowcell (Illumina).

### Low input HiC (LoHiC)

Between 0.5-1x10^6^ tumor cells were double crosslinked and subjected to HiC as previously described^56^. Then, single cells were processed with Chromium Next GEM single Cell ATAC Reagent Kits v2 (10X Genomics) following manufacture instructions.

### CRISPR/Cas9 screen

MM099 cells were transduced with lentivirus carrying a doxycycline inducible dCas9-KRAB-MeCP2 (Addgene, #140690) and a custom CRISPR library pRSGScribe1 (Cellecta) targeting 43 MES hubs plus 2 positive (*ACTB*, *GAPDH*) and 5 negative controls^57^ (5 sgRNAs/target) for a total of 230 sgRNAs. Single guide RNAs (sgRNAs) were designed using CRISPick^58^. After viral library titration, cells were infected with a 0.3 MOI to prevent multi-copy insertion and have a maximum of one sgRNA/cell. Finally, cells were collected at day 0 and 21 upon doxycycline induction (1μg/ml). Cell pellets were lysed to retrieve genomic DNA using QIAamp DNA Micro Kit (Qiagen, #56304). sgRNA sequencing libraries for t0 and t21 were prepared using NGS prep kit for sgRNA Libraries with supplementary primers (Cellecta, #LNGS-360, #LNGS-360-SP) following manufacturer’s instructions.

### Spectral partial wave spectroscopy (PWS) Imaging

PWS imaging was performed under physiological conditions (37°C, 5% CO_2_) using a stage-top incubation system (In Vivo Scientific). Cells were seeded on 8-chamber cover glass slides and allowed to adhere for 24 hours prior to imaging. Measurements were acquired on an inverted microscope (Leica DMIRB) equipped with a 63x oil-immersion objective (HCX PL APO, Leica), a CCD camera (ImageEM C9100-13, Hamamatsu), and a liquid crystal tunable filter (LCTF, Cri VariSpec) to enable spectrally resolved imaging. Broadband illumination was provided by a white-light LED source (X-Cite 120 LED, Excelitas), and a long-pass filter (Semrock BLP01-405R-25) was used to reject short-wavelength excitation light. Spectra image stacks were collected from 500 to 700 nm at 2-nm intervals, as previously described^59–61^.

PWS imaging quantifies wavelength-dependent interference patterns arising from multiple light scattering events within the sample, which are sensitive to nanoscale fluctuations in the local refractive index. Because refractive index is proportional to macromolecular density, PWS provides a label-free optical readout of chromatin nanoscale organization. For each pixel in the lateral image plane (x,y), the standard deviation of the backscattered intensity across the spectral dimension (𝜆) was computed, yielding a local chromatin packing scaling parameter, 𝐷_𝑎_(𝑥, 𝑦). The average nuclear chromatin packing scaling, 𝐷_𝑛_, was obtained by averaging 𝐷_𝑎_ across all pixels within the nuclear region of interest. 𝐷_𝑛_ reflects the organization of chromatin into spatially segregated packing domains and is derived from optical modeling and chromatin density autocorrelation functions established in prior chromatin tomography and PWS studies^61–63^. For each experimental condition, 𝐷_𝑛_ values were calculated for individual nuclei across biological replicates, and group-level statistics were used to quantify treatment-associated changes in chromatin packing.

In addition to live-cell measurements, PWS imaging was also performed on formalin-fixed paraffin-embedded tumor sections. OPAL-stained tumor sections were imaged by PWS and multi-channel fluorescence imaging. PWS and OPAL images were spatially registered, enabling chromatin packing metrics to be associated with immunophenotypically defined cell populations within intact tumor tissue.

### Dynamic PWS Measurements

Dynamic PWS imaging was performed as previously described^61, 64^ to quantify nanoscale chromatin motion in live cells. In this mode, wide-field backscattered intensity was recorded at a fixed wavelength (550 nm) over time, generating a three-dimensional data cube with dimensions (x, y, t). Temporal intensity fluctuations at each pixel arise from the movement of intracellular macromolecular structures, including chromatin. Fractional moving mass (FMM) was computed from the temporal variance of the backscattered intensity, 𝜎*_t_*^2^, and normalized using a biophysical scattering model to convert raw fluctuations into a quantitative measure of nanoscale mass motion. Following Gladstein *et. al.* (2019)^61^, FMM is defined as:

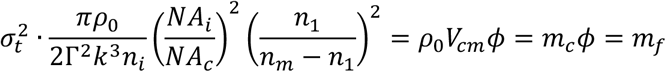

Where 𝑚_𝑓_ represents the mass undergoing motion within the sample and is given by the product of the mass of a typical moving cluster (𝑚_𝑐_) and the volume fraction of the mobile mass (𝜙). Here, 𝑚_𝑐_ = 𝜌_0_𝑉_𝑐𝑚_, with 𝑉_𝑐𝑚_ denoting the volume of a characteristic moving macromolecular cluster and 𝜌_0_ the dry mass density.

For normalization, the refractive index of a nucleosome was approximated as 𝑛_𝑚_ = 1.43, the nuclear refractive index as 𝑛_1_ = 1.37, and the refractive index of the immersion medium as 𝑛_𝑖_ = 1.518. The dry density of a nucleosome was taken as 𝜌_0_ = 0.55 𝑔 𝑐𝑚^−3^. The scalar wavenumber of the illumination light was 𝑘 = 1.57 × 10^5^ 𝑐𝑚^−1^, Γ denotes the Fresnel intensity coefficient for normal incidence, and the numerical apertures of collection and illumination were 𝑁𝐴_𝑐_ = 1.49 and 𝑁𝐴_𝑖_ = 0.52, respectively. As previously reported^61^, 𝜎*_t_*^2^ is sensitive to instrument-specific parameters, including depth of field and substrate refractive index. These effects were corrected using empirically determined calibration constants. Pixelwise FMM values were calculated for each nucleus, and distributions were compared across experimental conditions to quantify treatment-induced changes in chromatin mobility.

### sSMLM

Cells were seeded on 8-chamber cover glass slides (Cellvis, #C8-1.5H-N) and cultured to approximately 30% confluence. Samples were fixed in 4% paraformaldehyde prepared in phosphate-buffered saline (PBS) for 10 minutes at room temperature, followed by two PBS washes. After fixation, cells were quenched in 0.1% sodium borohydride in PBS for 7 minutes. After three PBS washes, cells were permeabilized and blocked in PBS containing 0.2% Triton X-100 and 3% bovine serum albumin (BSA) for 2hrs at room temperature. To label heterochromatic regions of chromatin, cells were incubated overnight at 4°C with anti-H3K9me3 antibody (Abcam, ab176916) at 1.2 µg/mL in blocking buffer. After 3 washes in PBS with 0.1% Triton X-100 and 0.2% BSA, cells were incubated with Alexa Fluor 647-conjugated secondary antibody (Thermo Scientific, #A-21245) at 2 µg/mL for 1 hour at room temperature. Cells were washed twice in PBS; to stain euchromatic regions, the labeling steps aforementioned were repeated using anti-H3K27Ac antibody (Thermo Scientific, #MA5-23516) and tagged with fluorophore Alexa Fluor 488-conjugated secondary antibody (Thermo Scientific, #A-11001) at 4 µg/mL in blocking buffer. To conduct imaging in STORM buffer containing 0.5 mg/mL glucose oxidase (Sigma-Aldrich), 40 µg/mL catalase (Roche), and 100 mg/mL glucose in TN buffer (50 mM Tris, pH 8.0, and 10 mM NaCl). Super-resolution imaging was performed using a Nikon Eclipse Ti-U inverted microscope equipped with a 100×/1.49 NA oil immersion objective (SR APO TIRF, Nikon) and an iXon Ultra 888 electron-multiplying CCD camera (Andor). A 637-nm laser (Obis, Coherent) provided excitation at the sample plane with an average power of 3–10 kW/cm² and using type F immersion oil (Nikon, #MXA22168). At least 10,000 frames were acquired per field of view with a 30 ms exposure time.

### Western blotting

Cells were harvested and lysed in LSDB buffer (0.5M KCl, 50mM Tris HCl pH 8.0, 20% Glycerol, 1% NP40, 1mM DTT) through three cycle of snap freezing in liquid nitrogen and thermobath at 37°C. Cell pellet debris was removed with the help of a pipette tip following centrifugation at 20.000g for 15min at 4°C. Between 10-20μg of proteins were run into pre-casted gels Nupage^TM^ 4-12% Bis-Tris (Thermo Scientific) using the dedicated apparatus (Thermo Scientific, #EI0001). Next, separated proteins were transfer into nitrocellulose membranes (Cytiva, #10600011) which were subsequently blocked into 5% milk TBST solution. Primary antibody staining was conducted by diluting antibodies in TBST (between 1:1000-5000 dilution, Supplemental reagents) and incubated overnight at 4°C with the corresponding membranes upon gentle mix. Secondary antibodies HRP-conjugated were diluted 1:10.000 in TBST and incubated 1hr at room temperature with gentle shaking. Protein signal was revealed using luminol substrate mix (Millipore, #WBLUR0500) and detected with an Odyssey XF Imaging System (LICORbio). Protein quantification was conducted using ImageJ.

### Immunoprecipitation (IP)

Protein immunoprecipitation was conducted as previously described^46^. Briefly, cells were harvested, pellet and gently lysed into 2 pellet volume of hypotonic buffer (10mM Tris HCl pH 7.6, 1.5mM MgCl_2_, 10mM KCl, 0.1% NP40) using mechanical shear with a loose Dounce to release the cytoplasmic fraction. Next, nuclei were pellet and resuspended in 1 volume of sucrose buffer (20mM Tris HCl PH 7.6, 15mM KCl, 60mM NaCl, 0.34M Sucrose) followed by a drop-by-drop gradual integration of 0.3 volumes of high salt buffer (20mM Tris HCl pH 7.6, 25% glycerol, 1.5mM MgCl_2_, 0.2mM EDTA, 0.9M NaCl) upon gentle vortex. After 30 min incubation at 4°C on constant slow spinning, ruptured nuclei were pellet, nuclear soluble fraction was collected and nuclei were further resuspended in 1 volume of sucrose buffer supplied with 1mM CaCl_2_ and 20U/μl of Micrococcal Nuclease (NEB, #M0247S), and incubated in thermoblock for 10min at 37°C. Micrococcal digestion was stopped by adding EDTA (final concentration 4mM) and lysate was centrifuged 20.000g for 30min at 4°C to obtain the nuclear insoluble fraction.

For CTCF co-IP, 500μg of nuclear insoluble fraction was diluted in 1mL of TGEN buffer (10mM Tris HCl pH 7.6, 150mM NaCl, 3mM MgCl_2_, 0.1mM EDTA, 10% glycerol, 0.01% NP40) for each IP. 1% of diluted extract was kept as western blot input. Next, between 5-10μg of primary antibodies against CTCF (Cell Signaling Technology, #2899), FOSL1 (Cell Signaling Technology, #5281), FOSL2 (Cell Signaling Technology, #19967), c-JUN (Cell Signaling Technology, #9165), pan-TEAD (Cell Signaling Technology, #13295) and IgG control (Cell Signaling Technology, #3900) were added in each IP and incubated overnight at 4°C on constant slow spinning. The day after, 25μl of magnetic beads A/G (Thermo Scientific, #88803) previously rinsed in PBS + 0.5% BSA were added to each IP and incubated 4hrs at 4°C upon slow spinning. Next, beads-antibody-proteins complexes were gently washed five times in cold TGEN buffer and eluted in LSDB and 1X Laemmli buffer in a thermoblock for 10min at 98°C before proceeding with western blot.

### Immunofluorescence (IF)

Tissue microarrays (TMAs) were sectioned at 5 µm sections and immunostained on a Leica BondRx auto-stainer according to the manufacturer’s instructions. In brief, sections were deparaffinized online and underwent antigen retrieval with either ER1 (Leica, AR9961; pH6) or ER2 (Leica, AR9640; pH9) retrieval buffer at 100o for either 20 minutes. Sections were then treated with 3% H2O2 to inhibit endogenous peroxidases. After blocking with Primary Antibody Diluent (Leica, AR93520), slides were incubated with the first primary antibody and secondary HRP polymer pair. Following secondary incubation, slides underwent HRP-mediated tyramide signal amplification with a specific Opal® fluorophore. Once the Opal® fluorophore was covalently linked to the antigen, primary and secondary antibodies were removed with a 95o retrieval step. This sequence was repeated 5 more times with subsequent primary and secondary antibody pairs, using a different Opal fluorophore for each primary antibody against human MITF (Sigma-Aldrich, #HPA003259), TCF4 (Abcam, #ab223073), S100B (Abcam, #ab115803), p-EGFR (Cell Signaling Technology, #4407), AXL (Cell Signaling Technology, #8661), and SOX10 (Sigma-Aldrich, #383R-15). After antibody staining, sections were counterstained with spectral DAPI (Akoya Biosciences, FP1490) and mounted with ProLong Gold Antifade (ThermoFisher Scientific, P36935).

Semi-automated image acquisition was performed on an Akoya Vectra Polaris (PhenoImagerHT) multispectral imaging system. Slides were scanned at 40X magnification using PhenoImagerHT 2.0 software in conjunction with Phenochart 2.0 and InForm 3.0 to generate unmixed whole slide qptiff scans. Image files were uploaded to the NYUGSoM’s OMERO Plus image data management system (Glencoe Software).

### Computational analysis

#### ChIPseq, CUT&Tag and ATACseq

Sequencing results were demultiplexed and converted in fastq using Illumina bcl2fastq software. Next, fastq files were trimmed and filtered according to the qc report obtained with TrimGalore (0.6.6, https://github.com/FelixKrueger/TrimGalore) and fastqc (0.11.7, https://www.bioinformatics.babraham.ac.uk/projects/fastqc/). For ChIP samples, fastq trimming was set for Illumina adaptors; for ATACseq and CUT&Tag, it was set for Nextera adaptors. Alignment to Hg38 human genome was performed using bowtie2 (2.4.1)^65^ and sam files were converted into bam using samtools (1.16)^66^. Bigwig files were generated from bam files using bamcoverage (deeptools 3.5.6)^67^ and filtered out for duplicate and bad genomic regions. Peak calling was performed using MACS2 (2.2.9.1)^68^ using paired-end mode, IgG (ChIP and CUT&Tag) was used as control, a q-value 0.05 threshold was set and both, narrowPeak and summit peaks were filtered for blacklisted regions using bedtools (2.31.0)^69^. An Hg38 CTCF whole genome track generated by JASPAR^70^ was used to filter CTCF peaks and maintain only the ones containing the CTCF motif. Peak annotation was performed with homer (4.10)^71^ using narrowPeak files and Hg38 genome parameters. DNA motif analysis of CTCF was done by first expanding of 200bp MACS2 summit peaks with bedtools slop, then fasta sequence was obtained using bedtools getfasta which were processed using MEME-ChIP^72^. DiffBind^73^ was used to identify state-specific peaks for CTCF, H3K27ac, ATAC and H3K9me3.

#### HiC

For HiC samples, fastq trimming, qc, alignment, valid pairs identification and matrix generation were performed using HiC-Pro workflow (3.1.0)^74^ as previously described^75^.

Briefly, valid pairs outputs were converted into juicer and cool formats using hicpro2juicebox and hicpro2higlass respectively (HiC-Pro). Matrix normalization was performed using cooler balance (cooltools)^76^ and addNorm (juicer tools)^77^. Copy number variations (CNVs) and structural variants (SVs) were detected using HiNT^78^ cnv and tl functions at 50kb BIN resolution and chromosomes’ segmentation were pooled into one file and converted in Log2 using python. Segments with a Log_2_≥0.5 and Log_2_≤-0.5 were considered amplified and deleted respectively. HiC-Pro valid pairs were filtered for CNVs and SVs using bedtools pairtobed. Compartment calling was conducted using eigs-cis function (cooltools) and 100kb BIN resolution matrixes. For compartment switch analysis, compartment score data from multiple cell lines were combined in a single text-delimited-file and delta switch was calculated as described in Narang et al^19^: compartments flipping sign with an absolute difference ≥0.5 and an absolute delta ≥1.2 were considered switched. Compartments’ saddle-plot was generated with cooltools. To measure H3K9me3, H3K27ac, ATAC and RNA RPKM enrichment in the compartments, bigwig files from these markers were segmented in 100KB BIN and averaged in each compartment using deeptools. Topological Associated Domains (TADs) identification was conducted using cooltools insulation providing an input matrix of 10kb BIN resolution, a 120kb window and Li threshold. From the tsv output, i) TADs boundaries were identified if considered ‘true’ under the ‘is_boundary’ column and ii) if the distance between two boundaries is less or equal to 3Mb was considered as a domain^79^. Loops were called using both 10kb and 5kb BIN matrixes with Peakachu^80^ and 10kb and 5kb were merged using hicMergeLoops (HiCexplorer). TADs boundaries and loops anchors co-occupied by CTCF were retained for downstream analysis using bedtools intersect and hicValidateLocation (HiCexplorer) respectively. HiC pile up maps, intra-TAD and inter-TAD contacts data were generated using GENOVA r package^81^. CTCF occupancy at loops anchors and TADs boundaries was calculated and visualized using deeptools. Loops and TADs were annotated using bedtools intersect and pairtobed against GENCODEv44 Hg38 bed file downloaded from UCSC Genome Browser. Graphs were generated combining Prism10, R and phyton.

#### ABC-score model

Active-by-Contact (ABC) model^82^ was generated following the authors guidelines (https://github.com/broadinstitute/ABC-Enhancer-Gene-Prediction) as previously described^75^. Briefly, cCRE were called and measured in activity using makeCandidateRegions and run.neighborhoods scripts in which ATACseq and H3K27ac bam and narrow.Peaks are integrated. Next, HiC matrix were processed using juicebox_dump and compute_powerlaw_fit_from_hic to extrapolate genome wide contacts. ABCscore for each active cCRE was measured using predict function in which cCRE, HiC contacts and RNAseq are integrated. Finally, cCRE were filtered using an ABCscore threshold of 0.02 and self-promoter contacts were excluded for downstream analysis. DESeq2^83^ was used to identify differentially active cCRE across melanoma cell lines.

To identify chromatin hubs, sumABC (Σ𝐴𝐵𝐶) was calculated for each cCRE as the sum of all contacted genes per cCRE and it was calculated with R (4.4.1) using the corresponding formulas:

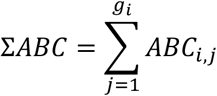

*ABC* = score

*g* = gene

Next, each cCRE was ranked based on the number of contacting genes with an expression level of Transcript Per Million (TPM) ≥1.

#### Polychrom

Chromatin 3D dynamics were simulated using a modified version of the Polychrom engine (OpenMM-based molecular dynamics, https://github.com/open2c/polychrom), implemented in our custom pipeline. Briefly, each chromosome is represented as a coarse-grained polymer, where individual 10 kb bins from Hi-C data correspond to “beads” connected by harmonic bonds (rest length = 1). Angular stiffness of each bead (*k*_θ_) is computed from integrated epigenetic, Hi-C, and TAD information:

- Epigenetic terms: local enrichment of H3K27ac and H3K9me3 (BigWig tracks)
- Compartment identity (A/B): sign(E1) from eigenvector decomposition
- TAD topology: TAD size-normalized scaling factor
- Hi-C contact environment: short-range, long-range, and inter-TAD interactions

Stiffness was modeled as:

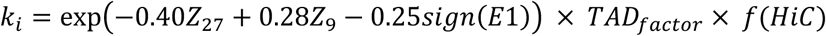

followed by clipping to a biophysically stable range (0.3–4.0). Softness is defined as:

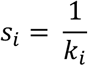

All forces used in the simulation include harmonic bonds, angular stiffness (per-bead *k*_θ_), excluded volume via polynomial repulsion and high-confidence Hi-C–derived long-range harmonic restraints (above a quantile threshold). An initial random-walk polymer is energy minimized and then propagated in Langevin dynamics for 150k integration steps (timestep 40 fs). Snapshots are saved every 5,000 steps using Polychrom’s HDF5Reporter, generating bead-track annotation, full 3D trajectory, contact map, per-frame radius of gyration (Rg) and curvature-based softness. Outputs include genome-wide stiffness tracks, compartment-stratified distributions, TAD-level summaries, and effective chromosome stiffness calculated from mean curvature across all trajectory frames. To quantify chromatin deformability under nuclear constriction, we implemented an additional energy-minimization pipeline. For each sample, the final equilibrium chromatin configuration from the Polychrom simulation is subjected to increasing levels of axial compression inside a narrowing slit. Given a slit half-height *h*, all beads satisfying |z| > *h* are energetically penalized by a soft-wall potential:

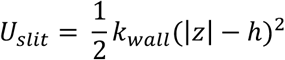

For each constriction level (typically *h* =1.0, 0.8, 0.6, 0.4, 0.3), the system undergoes NaN-safe local energy minimization and extraction of total system energy and principal radii of gyration (Rg1 ≤ Rg2 ≤ Rg3). This yields per-sample tables of Energy(h) and Rg(h). For each sample, deformability metrics were computed from the relaxed (largest slit size) and tight (smallest slit size) states:

- Constriction energy cost:

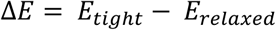

- Axial deformability (compression along Rg1):

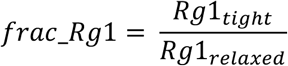

- Changes in chromatin anisotropy:

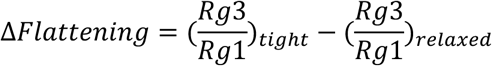

These metrics quantify whether chromatin becomes shorter, flatter, or more anisotropic under confinement. Statistical comparisons were performed in R using group-wise ANOVA and Kruskal– Wallis tests.

#### scRNAseq

PIPseeker v1.0.0 (Fluent Biosciences) was used to generate single cell matrix, barcode and features for each sample against Hg38 (human). Separate contig for GFP cDNAs was added to the genomes to detect malignant cells expressing a reporter. Cell calling, qc (including doublets removal) and clustering were done using Seurat5^84^. Melanoma cell states were assigned based on the score of previously established signatures^39, 85^ using AUCell^86^.

#### scATAC

The complete processing of fastq from alignment to single cell ATAC peak calling was performed using cellranger-atac-2.1.0 (https://github.com/10XGenomics/cellranger-atac). Next, scATACseq cellranger matrix was processed using Signac^87^ to analyze the overall ATAC distribution around TSS, fragments length, number of peaks per cell. Given that, cells with at least 100-3000 fragments, a blacklist ratio < 0.4, a nucleosome signal < 2 and a TSS enrichment > 1 were retained for downstream analysis. Malignant states were assigned to scATACseq object using Seurat5 FindTransferAnchors function and the scRNAseq matched object as a reference. Cell barcodes passing the qc and belonging to the assigned states were used to filter the paired fastq files which were then processed using the HiC analysis described above.

#### CRISPR/Cas9 screen

sgRNAs counts and dropout enrichment analysis comparing t0 with t21 were conducted using MAGeCK (v0.5.9.5)^88^.

#### RNAseq

Fastq were trimmed and evaluated for qc using TrimGalore (0.6.6) and fastqc (0.11.7). Then, trimmed fastq were aligned to Hg38 human genome and annotated using STAR (2.7.7a)^89^ using –quantMode TranscriptomeSAM GeneCounts and –sjdbGTFfile GENCODEv44. Using R (4.4.3), ReadsPerGene.out.tab were combined in a single table using data.table and processed for normalization and differential expression analysis using respectively edgeR^90^ and DESeq2^83^. Transcript per million (TPM) gene expression normalization was performed with rsem^91^ using Aligned.toTranscriptome.out.bam from STAR alignment as input.

#### Methylation

Raw IDAT files with 853,307 probes were imported and normalized in several consecutive steps using *SeSAMe* (version 1.12.9)^92^. A signal intensity-associated P-value was calculated based on out-of-band array hybridization (*pOOBAH*) and a cutoff of 0.05 was imposed. Background signal and dye-bias were corrected with *noob* and *dyeBiasCorr,* respectively. The methylation level of individual CpG sites was determined by calculating the beta (β) value, defined as the ratio of the fluorescent signal from the methylated allele to the sum of the fluorescent signals of both the methylated and unmethylated alleles. β values range from 0 (completely unmethylated) to 1.0 (fully methylated). β values were calculated using *getBetas.* Next, HM450 Hg38 probe manifest was downloaded from TCGAbiolinks^93^ and filtered for blacklisted regions. Genomic βvalues of each cell line were then inferred at bed regions of interests using GenomicRanges^94^. Analysis was conducted using R (4.4.1).

### Mouse experiments

12-273BM GFPLuc cell derived xenografts (CDXs) intended for loHiC were generated by injecting 1x10^4^ cells in 50% PBS + 50% Matrigel (Corning, #354234) intradermally into 8 weeks old NOD/SCID/IL2yR-/- male mice (RRID:IMSR_JAX:005557). Tumors were harvested once a ∼500mm^3^ volume was reached.

For metastatic dissemination studies, 1x10^5^ 501mel GFPLuc were resuspended in 100μl PBS and instilled by ultrasound-guided (Visualsonics Vevo 770 Ultrasound Imaging System) intracardiac injection into NOD/SCID/IL2yR-/- male mice 8-12 weeks old. Mouse distress and body weight were monitored over the course of the experiment until the established endpoint of ∼20% weight loss. After euthanasia, mice organs were dissected and imaged for GFP fluorescence on a Odissey M (LICORbio). GFP foci and signal intensity were quantified with ImageJ.

All mice were purchased from Jackson Laboratory and maintained in the NYULH animal housing according to the ethical compliance established by the NYU Institutional Animal Care and Use Committee (IACUC). All the experiments were approved and conducted under the IACUC protocol # IA16-00051.

### Data mining public datasets

TCGA SKCM transcriptomic data were downloaded from cBioPortal^95^, while SKCM HumanMethylation450 array data were obtained through Bioconductor package TCGAbiolinks. SKCM patients were classified as Melanocytic, Transitory, Neural-Crest and Undifferentiated based on a previous study^51^. Analysis was conducted using R (4.4.1).

DepMap Public 25Q2^96^ data was downloaded from the official website (https://depmap.org/portal). scRNAseq expression values from melanoma states in human^85^ were generated using a Seurat object retrieved from a public drive of the Marine Lab (https://rdr.kuleuven.be/dataset.xhtml?persistentId=doi%3A10.48804/GSAXBN).

ATACseq generated in TCGA tumors was retrieved from XenaBrowser^97^.

H3K27ac Hi-ChIP ValidPairs derived from TCGA tumors^98^ were downloaded from the NCI repository (https://gdc.cancer.gov/about-data/publications/TCGA-HiChIP-2024).

### Data and Code Availability

Supplementary data, raw data and the code to process the data generated in this work will be available in Gene Expression Omnibus and in a dedicated Github link after peer reviewed publication.

## Results

### Melanoma states exhibit distinct chromatin diffusion, density and domain maturation

At the nanoscale level (∼10nm), super-resolution imaging has revealed that chromatin is organized into CPDs, whose functions are determined by size (50-200nm) and packing scaling (D_n_), which we previously showed influence the probability of a cell to adopt a certain transcriptional state and nuclear physical properties^16, 62,99^. To investigate potential differences in CPDs across melanoma cell states as a measure of their nuclear physical properties, we performed high-resolution imaging on a panel of melanoma cell lines representing three main phenotypic states: melanocytic (MEL, n = 4), intermediate (INT, n = 3), mesenchymal (MES, n = 4), and of melanocytic cells upon MITF silencing (siMS, n = 2) vs. non-targeting control (siNTC) (**Fig. 1a**). Given that MITF directly regulates SOX10, siMS simultaneously abrogates the expression of both master regulators of melanocytic identity, and we tested whether their concomitant silencing elicits nuclear 3D changes. Partial Wave Spectroscopy (PWS) showed that MES cells exhibit lower CPD scaling (measured as D_n_) and more chromatin diffusion (measured as D_e_) than MEL cells, changes that were only partially recapitulated by siMS (**Fig. 1b**). As expected, INT cells displayed intermediate values between MEL and MES (**Fig. 1b**). To investigate if these differences were also observed in patient tissues, we integrated PWS with Opal multiplexed immunofluorescence (mIF) staining for melanoma cells (S100B+) in melanocytic (MITF+SOX10+) and mesenchymal (AXL, p-EGFR, TCF4) states in a patient tissue microarray (TMA) (**Fig. 1c**) containing cores representative of 20 unique primary melanoma cells (**Supplemental Table 1**). Following MEL and MES cell identification within the TMA (**Fig. 1d**, **Supp Fig. 1a**), melanoma cells classified as MES-like showed decreased CPD scaling compared to MEL-like cells (**Fig. 1e**) suggesting that CPD properties within melanoma cell states are also found in vivo.

**Figure 1:**
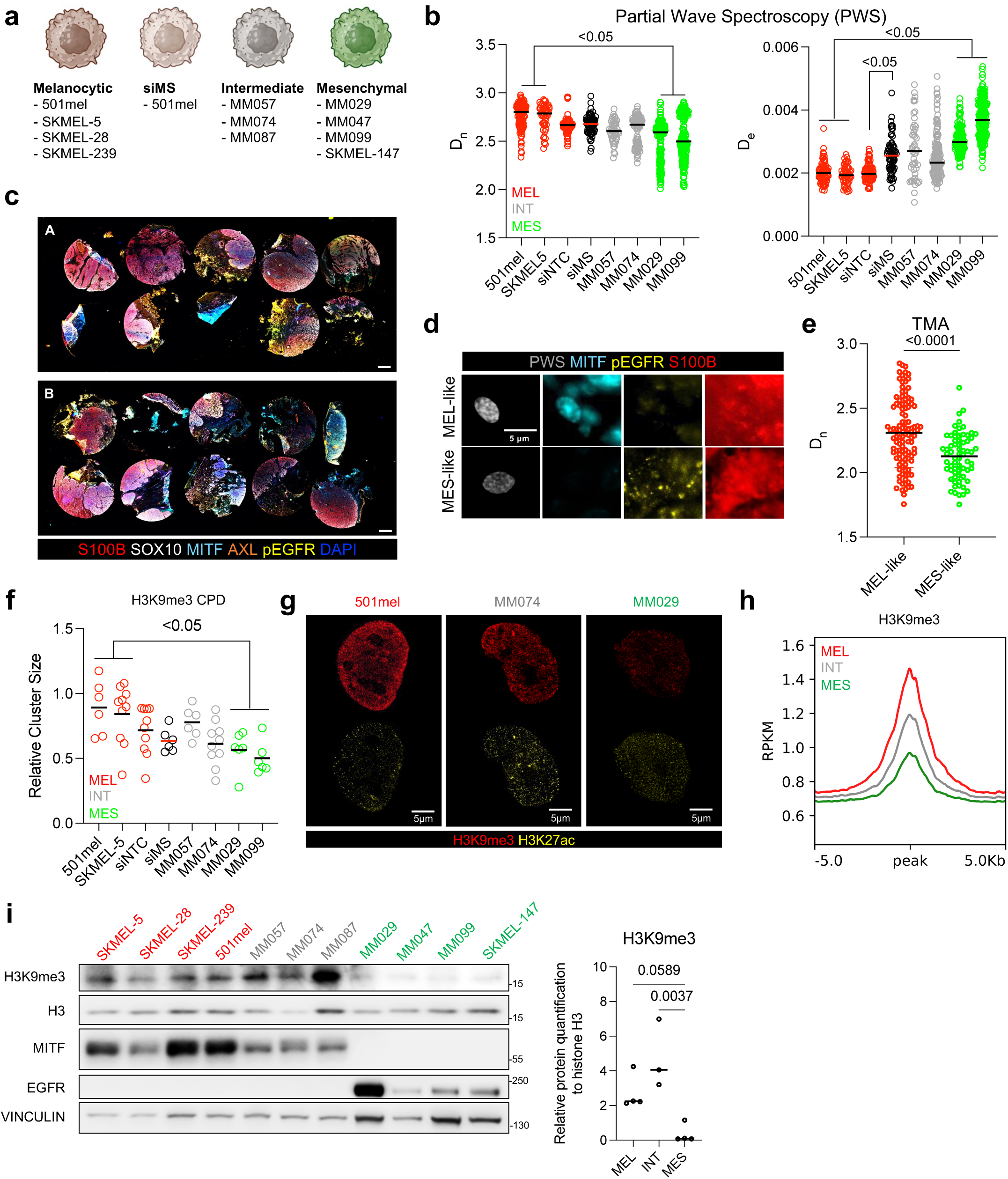
Melanoma cell states are characterized by distinguished chromatin nanoscale structures. **a**. Biorender illustration of the melanoma cell lines classified into states based on our and previous studies^45^. **b**. PWS quantification of Chromatin Packing Domain scaling (D_n_) and diffusion (D_e_) in a large panel of melanoma cell lines colored as melanocytic (MEL, red), intermediate (INT, grey) and mesenchymal (MES, green). Each dot represents a single cell, in black is shown the median bar. One-way ANOVA test p-values are shown and summarized as pairwise comparisons MEL vs MES. **c**. Whole slide snapshot of the Opal staining conducted in two TMAs (A-B) for S100B, SOX10, MITF, AXL and phospho-EGFR (pEGFR). Scale bars are referred to 5mm size. **d**. Representative MEL-like and MES-like cells identified in two melanoma TMAs by confocal microscopy using a combination of markers to measure PWS in melanoma cell states. **e**. Dot plot showing D_n_ measurements in MEL-like (n=109) and MES-like (n=67) cells in two melanoma TMAs. Each dot represents a single cell. Student unpaired t-test p-value comparison is shown. **f**. Relative H3K9me3 cluster size in different melanoma cell lines representative of MEL, INT, MES, siNTC and siMS groups. Each dot is the mean measurement in a single cell. One-way ANOVA test was conducted. Significant p-value summarizing MEL vs MES is shown. **g**. Representative sSMLM images for H3K9me3 and H3K27ac staining in three different melanoma cell lines colored based on their state of belonging (MEL = red, INT = grey, MES = green). **h**. H3K9me3 ChIP-seq global metaprofile in MEL, INT and MES cells. Each profile is the average of at least three biological replicates (different cell lines). **i**. (Left) Immunoblot for different proteins in a large panel of melanoma cell lines marked in red (MEL), grey (INT) and green (MES). MITF and EGFR are enriched in MEL/INT and MES cells respectively. (Right) H3K9me3 relative quantification to total histone H3 based on the blot on the left. Unpaired t-test was performed for pairwise comparisons. P-values are displayed.

Heterochromatin and euchromatin have been previously linked to higher and lower mechanical stiffness, respectively^100^. CPD maturation relies on H3K9me3 heterochromatin accelerating packing in the CPD core, while euchromatin, characterized by higher H3K27ac, stabilizes the peripheral CPD zone^101^. To gain more insight into CPD maturation related to cell states, we performed sSMLM for H3K9me3 and H3K27ac in our panel of cell lines. In line with our hypothesis, CPD maturation is reduced in MES cells, as indicated by a significant decrease in H3K9me3-positive CPD size and number relative to MEL cells (**Fig. 1f-g**, **Supp Fig. 1b**), findings further supported by a global genome-wide loss of H3K9me3 by ChIP-seq (**Fig. 1h**). No significant differences were observed in H3K27ac by sSMLM (**Fig. 1g**, **Supp Fig.1b**), ChIP-seq (**Supp Fig. 1c**) nor upon siMS (**Supp Fig. 1d-e**).

Altogether, we demonstrate that MES cells are characterized by increased chromatin diffusion and reduced chromatin density, explained by a reduction in CPD number, size, and maturation, as well as global loss of the H3K9me3 mark, suggesting decreased nuclear stiffness. These differences are only partially mimicked by MITF and SOX10 silencing (siMS), suggesting that additional mechanisms are required to fully recapitulate the nuclear topology and biophysical properties of MES cells during phenotype switching.

### Inactive compartments and heterochromatin are reshaped during melanoma phenotypic switch

The genome has two major, micron-sized spatial compartments, A and B, associated with active and inactive chromatin respectively^102, 103^ within which CPDs are embedded ^16^. Using Cooltools^76^, we examined melanoma cell lines by sub-chromosomal compartments A and B, further validated by integrating H3K27ac and H3K9me3 ChIPseq, ATACseq, and RNAseq in our panel of cell lines. As expected, A-type compartments are enriched for H3K27ac, ATAC, and RNA signal compared to B compartments, which instead are enriched in H3K9me3 (**Supp Fig. 2a**). To identify state-specific changes in A and B compartments, we performed one minus cosine similarity Kmeans clustering and found a subset of A compartments specific to each phenotype (MEL and MES; **Fig. 2a**). Although no change in the global proportion of A and B compartments was observed across states or upon siMS (**Supp. Fig. 2b**), saddle *cis* compartment quantification shows an increase in B-to-B interactions in INT, MES, and siMS cells relative to MEL and siNTC, respectively (**Fig. 2b-c**). No significant differences were found across *trans* compartments interactions (**Supp Fig. 2c-d).** We analyzed compartment switching from A-to-B and B-to-A between states using a recently published model^19^. Consistent with a switch, epigenetic and transcriptomics marks were found significantly changed from A-to-B and B-to-A comparing MEL, INT and MES, but not in siMS (**Supp Fig. 2e**). We found < 400 compartment switches between MEL and INT, while almost 2000 compartments switched between MEL and MES, predominantly from B-to-A which is triggered as well by siMS (**Fig. 2d**). This is in line with the observed decrease in H3K9me3 (**Fig. 1h-i**) and CPDs maturation by sSMLM (**Fig. 1f-g**). In line with our previous observation of decreased H3K9me3 from MEL to MES, our 3D chromatin data indicate that the melanoma phenotypic transition from MEL to INT and MES is marked by a B-to-A compartment switch, mimicked by siMS.

**Figure 2:**
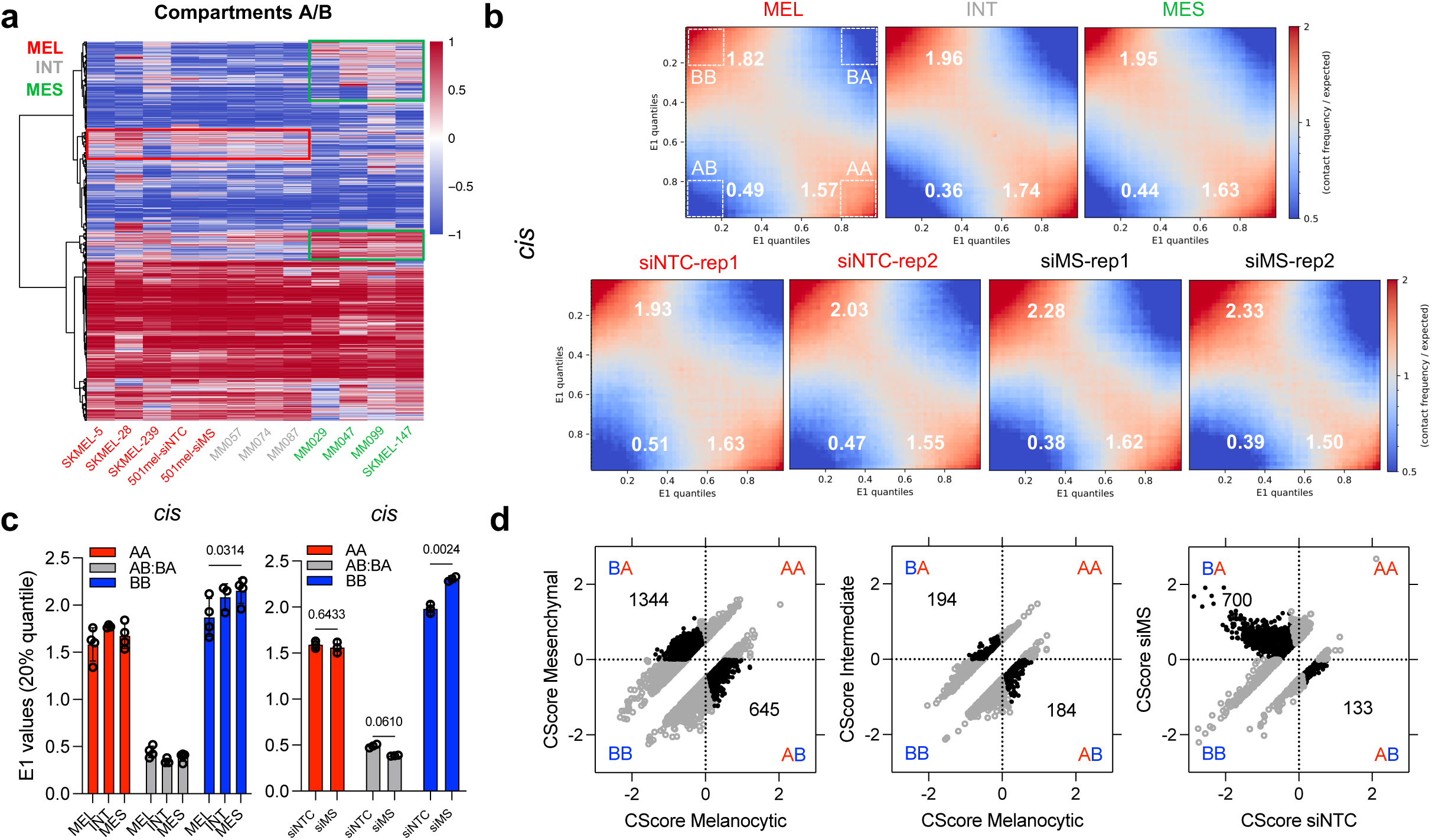
Nuclear compartments switch occurs during phenotype switching. **a.** Whole genome *cis* compartments A and B clustering in melanoma cell lines. In red and green are marked A compartments specific of melanocytic and mesenchymal states respectively. Color scale represents compartment score z-score normalized by column. Ward hierarchical clustering dendrogram is shown on the left. **b.** States, siNTC and siMS *cis* compartments saddle plots. 20% quantile areas at the corners were measured (as marked by dotted white squared) as representative of more radical interactions across A and B. **c.** Group histogram of *cis* AA, BB, and AB:BA interactions in states (left), siNTC and siMS (right). Two-way ANOVA test was used, p-values are displayed. **d.** Compartment switch analysis from BA, AB, AA and BB in melanoma states, siNTC and siMS. Significant switches are shown in black, the number of switch events are displayed in the corresponding areas.

### MES cells have a reduced number of topologically associated domains

To further interrogate 3D chromatin changes associated to cell states, we examined the number, size and distribution of TADs (contact hubs of genes, promoters, and enhancers) and quantified insulation scores. We found MES cell lines have significantly fewer (**Fig. 3a***, top*), larger (**Fig. 3a***, bottom*) TADs than MEL and INT cell lines. However, MITF and SOX silencing (siMS) did not impact TAD size or number (**Fig. 3a**). Next, TAD activity was evaluated by first averaging the TAD boundary strength, measured by insulation score. MES cells showed an overall decrease in insulation and average CTCF occupancy at MEL and INT TADs boundaries (**Fig. 3b, Supp. Fig. 3a-b**), whereas siMS display only decreased CTCF binding at boundaries but no change in insulation (**Supp. Fig. 3c**). TAD contact frequency was measured using Aggregate Topological Analysis (ATA), which revealed a global increase of contacts in the inter-TAD surroundings in INT and MES cells (**Fig. 3c**). In parallel, we observed significantly more inter-TAD contacts between +1, +2 and +3 neighbor TADs in INT and MES cells than in MEL cells (**Fig. 3d**). siMS had no effect on intra and inter-TADs contacts (**Supp. Fig. 3d-e**).

**Figure 3:**
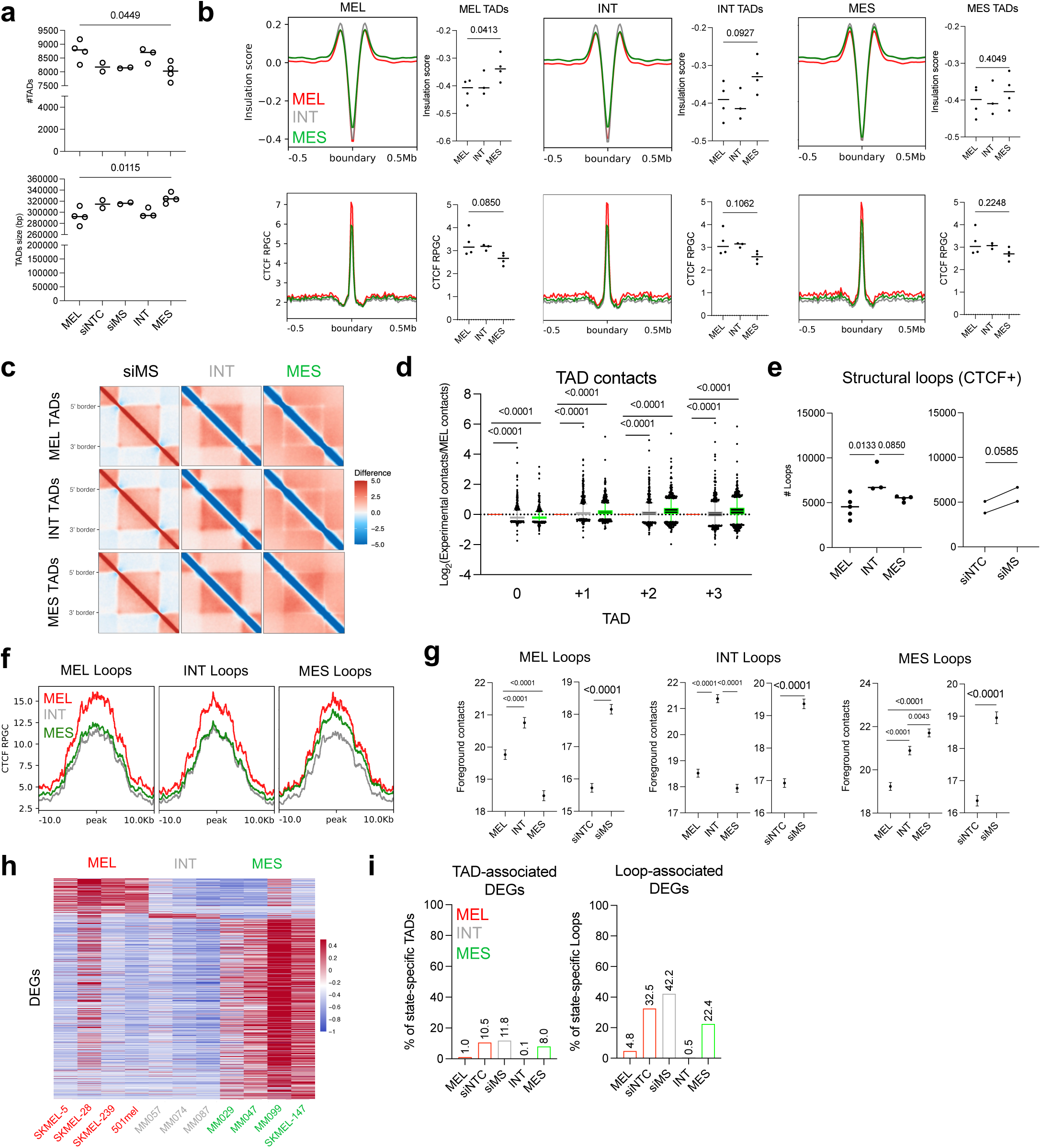
Structural domains and loops properties of melanoma cell states. **a.** Column dot plot of the number (top) and size (bottom) of TADs identified in melanoma states, siNTC and siMS. One-way ANOVA significant p-values are shown. Each dot represent a biological replicate. **b.** Insulation score (top) and CTCF binding (bottom) profiles of melanoma states compared at melanocytic, intermediate and mesenchymal TAD boundaries. Summit values are quantified and statistically compared in the sided column dot plots. One-way ANOVA test p-values are displayed. Each dot is representative of a biological replicate. **c.** Aggregate TAD analysis heatmap showing the relative difference in contacts of MES, INT and siMS cells against MEL/siNTC at melanocytic, intermediate and mesenchymal TADs. **d.** Bar dot plot quantifying the *cis* interactions in INT and MES cells relative to MEL cells as intra-TADs (0) and inter-TADs between neighbor TADs (+1, +2, +3). One-way ANOVA test was conducted independently in each TAD group (0, +1, +2, +3). P-values are shown. Each dot represent an independent interaction. **e.** Structural loop quantification in melanoma states (left), siNTC and siMS (right). One-way ANOVA and paired t-test were conducted for left and right plots respectively. Dots represent biological replicates. **f.** CTCF mean genomic occupancy at melanocytic, intermediate and mesenchymal loops’ anchors in melanoma states. **g.** HiC contacts frequency at loops anchors (foreground) at melanocytic, intermediate and mesenchymal loops relative to the background (Foreground contacts) in melanoma states, siNTC and siMS. One-way and paired t-test analysis were performed for melanoma states, and siNTC/siMS plots respectively. P-values are shown. **h.** Differentially expressed genes (DEGs) heatmap identified in melanoma states. Color bar represents z-score of TPM expression normalized by row. **i**. Bar plot quantification of the percentage of TADs (left) and loops (right) associated with DEGs that overlap with state-specific TADs and loops respectively.

Next, we identified loops and filtered for structural loops occupied by CTCF at both anchors. Notably, INT cells have the highest number of loops compared to other states, while siMS significantly increased the number of loops relative to siNTC (**Fig. 3e**). Similarly to TADs’ boundaries, CTCF occupancy at loops’ anchors is reduced in MES, INT and siMS cells (**Fig. 3f, Supp Fig. 3f**). Aggregate Peak Analysis (APA) shows an overall enrichment of contacts at the loops’ foreground (#contacts foreground - #contacts background) of the corresponding state, except for INT cells that have higher activity, mimicked by siMS cells compared to siNTC (**Fig. 3g, Supp. Fig. 3g**).

Finally, we intersected boundaries and loop anchors to identify unique TADs and loops across states and in siNTC vs siMS (**Supp. Fig. 3h**), and cross-referenced them with differentially expressed genes (DEGs) identified by RNAseq (**Fig. 3h**, **Supp. Fig. 3i, Supplemental Table 2**). Remarkably, only a modest fraction of state-specific TADs were found associated to DEGs for all states and siMS; i.e., only 1-10% of state-specific TADs result in DEGs (**Fig. 3i**, *left*). Similarly, structural loops were poorly associated to DEGs; except for siNTC, siMS and MEL, with ∼20-30% of specific loops resulting in DEGs (**Fig. 3i**, *right*). This indicates that differences in TAD and loop activity between states are disproportional to the changes in gene expression—particularly in MES cells, suggesting the possibility that changes in chromatin structure during the MES-to-MEL phenotypic transition are not just responsible for gene expression changes but may confer some other advantage to MEL cells.

In sum, melanoma dedifferentiation from MEL to MES is associated with enlarged domains and increased inter-TAD communications, in line with potentially enhanced nuclear and chromatin flexibility. Intermediate (INT) cells present multi-state loops and thus an overall higher number of loops, suggesting a ‘poised’ 3D chromatin state, which is in part recapitulated by siMS.

### State-specific chromatin architecture features are conserved *in vivo*

We have shown that melanoma phenotypic states display specific 3D chromatin architectures that correlate with nuclear physical properties, irrespective of mutant drivers (i.e., BRAF, NRAS). To complement these *in vitro* findings, we asked if melanoma states adopt similar state-specific 3D chromatin architecture *in vivo* using: i) scRNAseq combined with loHiC and ii) bulk HiC on chromatin from FFPE specimens from melanoma patients. We performed loHiC on a cell-derived xenograft (CDX) from a patient-derived melanoma brain metastasis short term culture (12-273BM) ^104, 105^, which was able to recapitulate the intra-tumor heterogeneity observed in clinical specimens. MEL and MES states were identified by scRNAseq through Seurat and integrated into the loHiC Signac clusters using Seurat bridge integration^84^ (**Fig. 4a, Supp. Fig. 4a**). Next, we performed pseudo-bulk HiC analysis on the MEL and MES clusters for which we obtained a good, genome-wide HiC contact coverage (**Supp. Fig. 4b**), and confirmed their malignant status using copy number variation analysis (**Supp. Fig. 4c**). Remarkably, we validated most of our in vitro observations, such as increase in BB compartment interactions (**Fig. 4b, Supp. Fig. 4d**), fewer and larger TADs (**Fig. 4c**), weaker insulation (**Fig. 4d**) and decreased intra-TAD contacts in MES vs MEL cell states (**Supp. Fig. 4e**). Similarly, bulk HiC in a small cohort of patients’ FFPE tumor samples broadly classified as MEL or MES based on histopathological characteristics (**Supplemental Table 3**) showed a trend to a reduction in the number of TADs in MES tumors (**Fig. 4e**). Overall, these data indicate that state-specific 3D chromatin features are not an artifact of in vitro culture but an actual feature of melanoma tumors.

**Figure 4:**
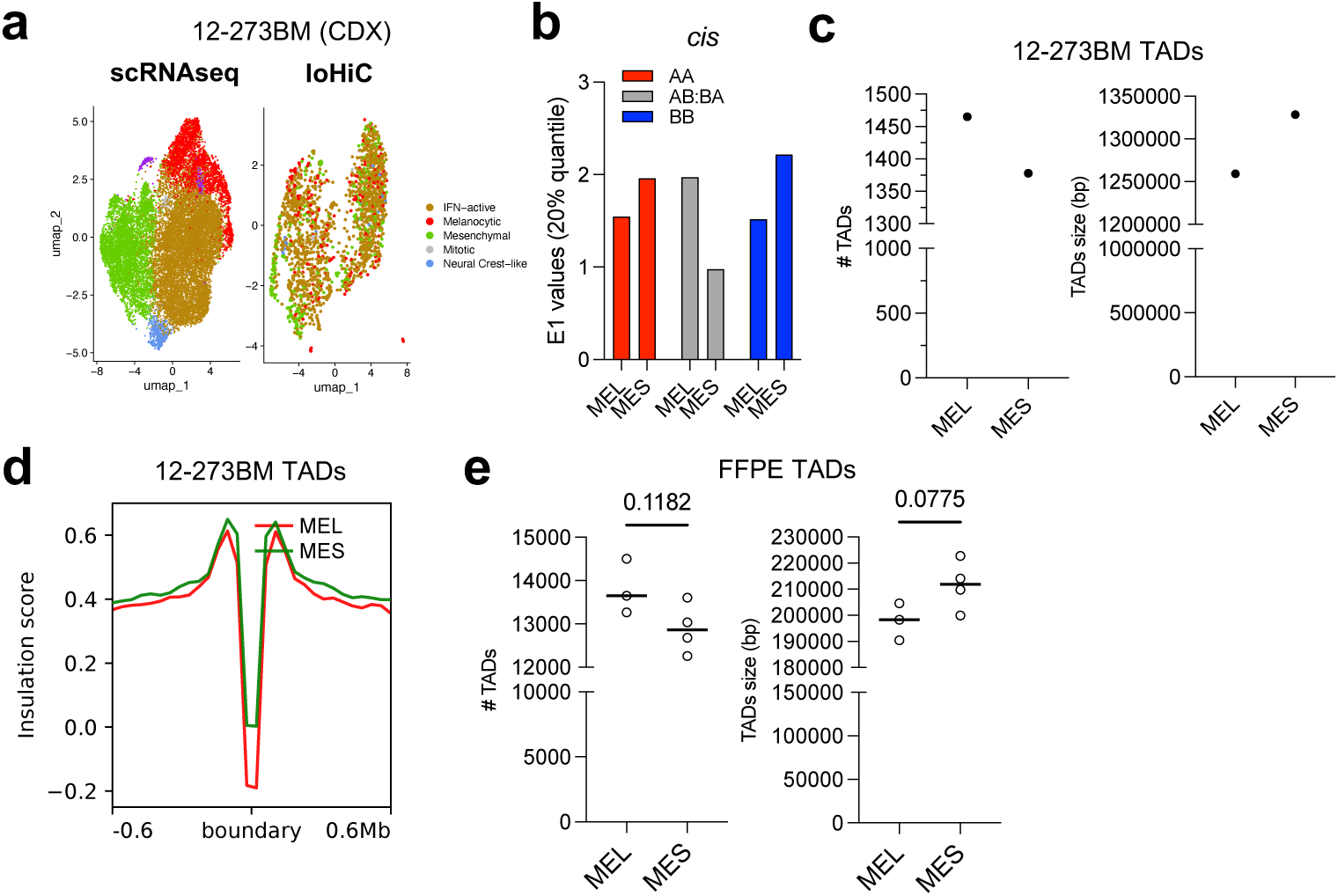
Melanoma states 3D chromatin differences are found *in vivo*. **a.** Seurat UMAP representing the bridge integration of melanoma cell states between scRNAseq (left) and loHiC (right) obtained from a cell derived xenograft (CDX) of 12-273BM short-term patients’ culture. **b.** cis compartment interactions quantified in MEL and MES states identified in CDX tumors. **c.** Column dot plot showing the quantification of the number (left) and size (right) of TADs in MEL and MES states by loHiC. **d.** Genome-wide insulation score profile at TADs’ boundaries in loHiC MEL and MES states. **e.** TADs’ number (left) and size (right) quantified in a small cohort of FFPE MEL (n=3) and MES (n=4) tumors processed with HiC. Unpaired t-test was conducted and resulted p-values are shown. Each dot represent a patient sample.

### Compression during migration induces a MES-like rewiring of nuclear architecture in MEL cells, suppresses heterochromatin levels, and enhances their metastatic potential

While mechanical forces are known to modulate chromatin accessibility^31, 32,33^ and viceversa^25, 100, 106–108^, the reciprocal influence of 3D chromatin conformation on nuclear flexibility in the context of phenotype switching is poorly understood. We have shown that MES cells, compared to MEL cells, possess a more ‘relaxed’ nuclear architecture, characterized by a lower number and less mature of CPDs **(Fig. 1f-g**), B-to-A compartment switching (**Fig. 2d**), fewer number of TADs with weaker insulation (**Fig. 3a-b**), and reduced heterochromatin (**Fig. 1g-i**). We hypothesized that this ‘loose’ nuclear conformation allows MES cells to better withstand the mechanical stress of invasion or intra/extravasation, and that reduced physical constriction can in turn remodel the 3D genome towards a pro-metastatic state.

To determine this, we integrated HiC, H3K27ac, and H3K9me3 profiles to model chromosome 1 chromatin as a Hamiltonian coarse-grained polymer bead chain using Polychrom (Open2C). Here, we simulate angle bending upon artificial scaled compression forces to measure stiffness and cost of motion of the polymer. The radius of gyration (Rg) measures the extension and compaction capacity: stiffer regions will adopt extended and rigid conformations increasing Rg, while softer regions can collapse reducing Rg and additionally facilitate *cis*-long interactions (**Supp. Fig. 5a**). In line with our hypothesis, our simulation predicts that MES cells have lower TAD and chromosome stiffness (**Fig. 5a),** as well as reduced energy cost upon simulated constriction and a trend to increased deformability (**Fig. 5b**, **Supp. Fig. 5b**). To experimentally prove this finding, MEL and MES cells were seeded in 3μm-pores transwells for 72hrs in complete media to allow cells to migrate and constrict spontaneously. Next, both non-migrated (Pre = on top side of the transwell) and migrated (Post = on lower side of the transwell) cells were processed for HiC, RNAseq, CUT&Tag (H3K27ac, CTCF and H3K9me3), and ATACseq. Post-migration, MEL cells were associated with gained MES-like features such as lower melanocytic gene expression (i.e., MITF, SOX10; **Fig. 5c, Supp. Fig. 5c**) reduced H3K9me3 levels (**Fig. 5c-d**), increased BB compartments interactions (**Fig. 5e, Supp. Fig. 5d-e**), fewer TADs (**Fig. 5f**), and decreased insulation and CTCF binding at TAD boundaries (**Fig. 5g** In contrast, chromatin and gene expression of MES cells were only modestly affected (i.e., EGFR, AXL; **Fig. 5c**), except for an increased melanoma mesenchymal signature, as if migration ‘exacerbates’ MES cells identity (**Supp. Fig. 5f**).

**Figure 5:**
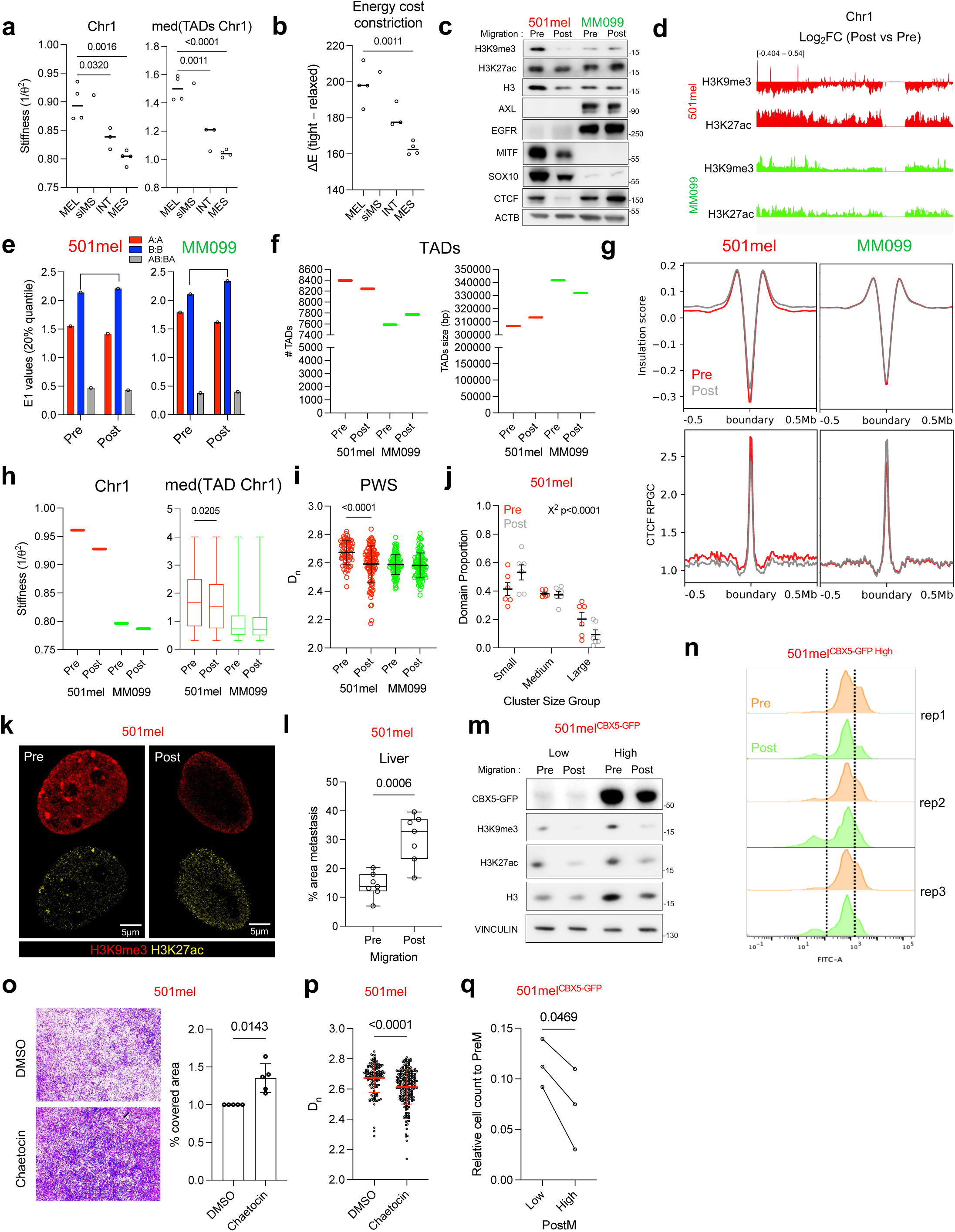
Nuclear constriction actively rewire 3D chromatin in a state-specific manner. **a.** Column dot plot quantification of chromatin stiffness predicted by polychorm in chromosome 1 (left) and the median stiffness per TAD of chromosome 1 (right) in melanoma states and siMS. P-values obtained from one-way ANOVA study are shown. Each dot is a biological replicate. **b.** Predicted energy cost of chromosome 1 subjected to a simulated constriction gradient. One-way ANOVA test was conducted for statistical comparison. Significant p-values are shown. **c.** Representative immunoblot for multiple targets detected in whole cell protein extracts from 501mel (MEL) and MM099 (MES) Pre and Post migration. ACTB and H3 serves as loading controls. Relative molecular weights are shown on the right. **d.** Bigwig comparison between Pre and Post genomic occupancy at chromosome 1 of H3K9me3 and H3K27ac in 501mel and MM099. Tracks are equally scaled spanning a range displayed on top left. **e.** Grouped histogram quantification of the 20% quantile *cis* compartments AA, BB, AB:BA interactions in 501mel and MM099 Pre and Post migration. Brackets are shown to highlight BB comparison between Pre and Post. **f.** Quantification of the number (left) and size (right) of TADs in 501mel and MM099, Pre and Post migration. **g.** Insulation score (top) and CTCF occupancy (bottom) profiled in Pre and Post cells at TADs boundaries of 501mel and MM099 cells. **h.** Stiffness quantification of whole chromosome 1 (left) and median TAD stiffness of chromosome 1 (right) in Pre and Post 501mel and MM099 cells. One-way ANOVA test was conducted. **i.** D_n_ measurement by PWS in 501mel and MM099 Pre and Post cells. Unpaired t-test comparisons between Pre and Post were conducted and significant p-value is shown. Each dot represents a single cell considered as biological replicate. **j.** H3K9me3 domain proportion in 501mel Pre and Post cells. Based on their radii size, domains were classified in small (≤0.55nm), medium (55-80nm) and large (>80nm). Each dot represents the average of independent measurements in single cells. Chi-square test of independence was conducted. Summary p-value is shown. **k.** Representative images captured by sSMLM for H3K9me3 and H3K27ac in 501mel Pre and Post migration. **l.** Box dot plot quantification of liver metastasis area (%) derived from NSG animals injected systemically (intracardiac) with 501mel Pre and Post. Each dot represents a biological replicate. Unpaired t-test was performed and p-value is shown. **m.** Immunoblot comparing Pre and Post 501mel^CBX^^5-GFP^ whole cell protein extracts carrying low or high levels of CBX5-GFP. Protein targets and relative molecular weights are shown at left and right respectively. Vinculin and H3 serve as loading controls. **n.** CBX5-GFP quantification by flow cytometry (FITC-A) in Pre and Post migration 501mel CBX5-GFP High. Dotted lines serve as visual reference to appreciate the appearance of low and decrease of high CBX5-GFP cells after migration. Three biological replicates are shown. **o.** (Left) Representative image of post migration cells upon DMSO or Chaetocin. (Right) Bar dot plot quantification of 501mel migratory cells upon DMSO or Chaetocin (5nM) based on crystal violet staining of transwell. Paired t-test p-value is shown. Each dot represent a biological replicate, and each replicate is the average of three technical replicates. **p.** Dot plot quantification of D_n_ measured by PWS in 501mel cells treated with DMSO (control) or Chaetocin. Each dot represent a single cell. Mean and standard deviations are shown in red. Unpaired t-test was performed, p-value is shown. **q.** Relative cell number of post migratory cells to pre migration of 501mel^CBX^^5-GFP^ High and Low. Each dot is a biological replicate. Paired t-test was conducted, p-value is displayed.

Polychrom simulation in Pre and Post cells predicted a significant decrease in stiffness in Post MEL relative to Pre MEL, but not in Post MES cells relative to its Pre counterpart (**Fig. 5h**). Consistent with this finding, CPD scaling (D_n_) measured by PWS was significantly reduced in Post cells, particularly in MEL cells (**Fig. 5i**, **Supp Fig. 5g**). In parallel, MEL Post cells displayed a significant reduction in mature large H3K9me3 CPDs and a concomitant increase in small ones (**Fig. 5j**) together with a global decrease in H3K9me3 by sSMLM (**Fig. 5k**) which were not observed in MES Post cells (**Supp Fig. 5h**). While constriction did not change their proliferative capacity (**Supp. Fig. 5i**), Post MEL cells exhibited increased metastatic potential to the liver upon intracardiac injection relative to their Pre counterparts (**Fig. 5l, Supp. Fig. 5j**).

Next, we tested whether Post cells were a product of selection of a pre-existing subpopulation in the initial culture, or whether physical constriction during migration actively results in cells with Post features. To do that, we infected MEL cells with a doxycycline inducible CBX5-GFP construct at 0.3 MOI (∼ 0.3 plasmid per cell) as a reporter of H3K9me3 content within the cells. Confirming the value of this construct as a marker of heterochromatin, we observed increased H3K9me3 levels in FACS sorted CBX5-GFP^High^ cells compared to CBX5-GFP^Low^, and a decrease of these markers upon post migration (**Fig. 5m**). Flow cytometry on Pre and Post CBX5-GFP^High^ cells showed the emergence of a CBX5-GFP^Low^ subpopulation Post migration (**Fig. 5n**). These data suggest that physical confinement during migration does not passively select but *actively* triggers a H3K9me3 low subpopulation.

We further tested whether reducing H3K9me3 global content in MEL cells is sufficient to increase nuclear flexibility and cell migration observed in MES cells. Only 12-24hrs of treatment with the histone methyltransferase (HMT) SUV39H1/2 inhibitor, Chaetocin increased cell migration (**Fig. 5o, Supp Fig. 5k**) and reduced H3K9me3 without affecting the levels of melanocytic markers such as MITF and SOX10 (**Supp Fig. 5l**). In line with our hypothesis of heterochromatin being a physical barrier to migratory constriction, a short pulse of Chaetocin (24hrs) was sufficient to significantly decrease D_n_ as a proxy of nuclear stiffness (**Fig. 5p**), and sorted CBX5-GFP^Low^ (low H3K9me3) cells displayed higher migratory capacity than CBX5-GFP^High^ (high H3K9me3) cells relative to their non-migrated counterpart (PreM) (**Fig. 5q**).

These data indicate that MEL cells gain MES features as they intravasate and that treatments that make MEL 3D chromatin ressemble the MES state are sufficient to confer increased migratory potential before changes in gene expression are observed.

### CTCF genomic occupancy reveals a transcriptional activator function specific to mesenchymal cells

We observed that MES cells have a global reduction of CTCF at TAD boundaries compared to MEL cells (**Fig. 3b**), which is in line with an overall CTCF protein decrease in MES (**Supp. Fig. 6a**). However, mean CTCF occupancy at TSSs is higher in MES compared to MEL and INT cells (**Fig. 6a**), suggesting a CTCF relocation from insulators to regulatory elements. To test this hypothesis, we first identified all active candidate *cis* regulatory elements (cCREs) by integrating ATACseq, H3K27ac ChIPseq, RNAseq and HiC using the Activity-by-Contact (ABC) model^82^ in all cell lines. Next, we intersected CTCF peaks with the ABCmodel identified cCREs^82^ to define the proportion of CTCF binding at structural-like (ABC-) vs. enhancer–promoter (ABC+) regions. This analysis demonstrated a global increase of CTCF binding to cCRE in MES and siMS cells compared to MEL cells (**Fig. 6b**). In line with the reduced insulator function in MES cells, HiC pileup map analysis at CTCF peaks shows a strong increase in global contacts as well as between adjacent 5’ and 3’ regions (span in graph) in MES cells, indicating a reduced level of conventional insulation activity compared to MEL, INT and siMS cells (**Fig. 6c-d, Supp. Fig. 6b**). Conversely, differential binding analysis of CTCF between MEL and MES cells (**Supp. Fig. 6c**) showed increased occupancy of MES CTCF at cCREs relative to structural-like (CTCF-only) anchors (**Fig. 6e**). These MES-specific CTCF bound regions are co-occupied by MES TFs such as TEAD4 and FOSL2 (**Fig. 6f**) and associated mostly with enhancers (n = 1,479 ABC cCREs) (**Supp. Fig. 6d**), suggesting that CTCF may exert a direct gene regulatory function in concert with AP1/TEADs in MES cells. Accordingly, CTCF and AP1/TEADs co-immunoprecipitated (**Fig. 6g**), further supporting these TFs binding as part of CTCF complexes. Moreover, TF motif footprint analysis demonstrates a significant increase in CTCF motif accessibility in MES and siMS compared to MEL and siNTC (MEL vs MES, Lg_2_FC = -0.35251; siNTC vs siMS, Lg_2_FC = -0.40685) (**Fig. 6h**). To identify CTCF regulated genes in MES cells, we looked at the activity and genes directly contacted by the ABC cCREs overlapping with CTCF MES peaks (**Supp Fig. 6e**), finding 5,081 cCRE-associated genes (**Supplemental Table 4**). We confirmed that putative CTCF-regulated MES genes are upregulated in MES cell lines^51^ and patient samples^85^ relative to MEL cells (**Supp. Fig. 6f-g**). To determine if MES genes depend on CTCF, we performed a brief knockdown of CTCF (24hrs) using siRNA followed by RNA-seq in MM099 (MES) (**Fig. 6i**, **Supp. Fig. 6h**). Notably, we found a significant overlap (n = 852) between CTCF ABC-predicted contacted genes (n= 5,081) and siRNA differentially expressed genes (n = 4,037) (**Supp. Fig. 6i**). Strikingly, CTCF depletion leads to a significant increase in MEL and decrease in MES gene signatures^53^ (**Fig. 6j**), and siCTCF upregulated and downregulated genes are found significantly associated to MEL and MES states respectively in patients (**Supp. Fig. 6j**), suggesting that CTCF is indeed required to maintain MES identity.

**Figure 6:**
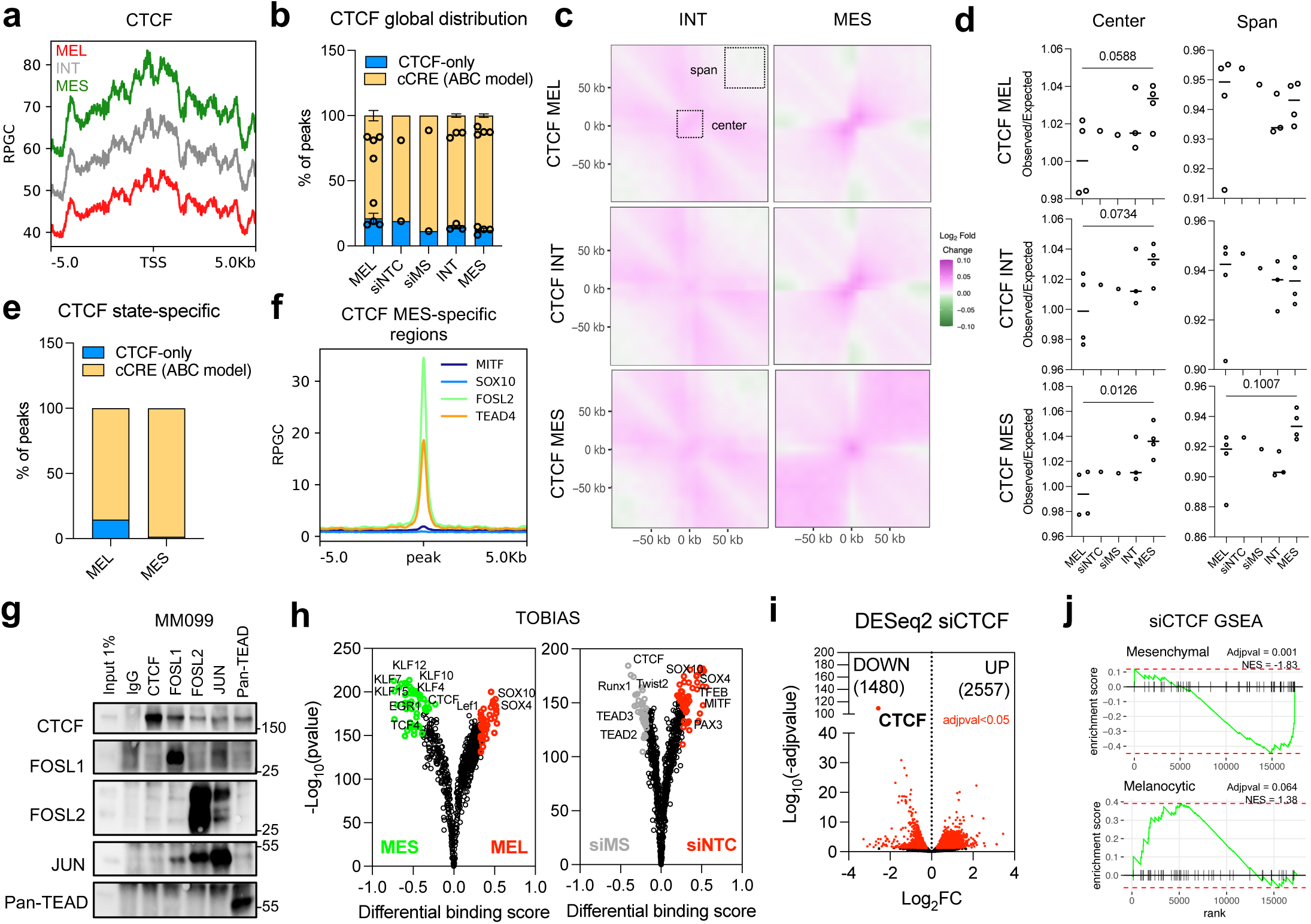
CTCF is a mesenchymal identity transcriptional regulator. **a.** Metaprofile occupancy of CTCF at transcription start sites (TSS) in MEL, INT and MES cells. **b.** Grouped bar plot representing CTCF occupancy at candidate *cis* regulatory elements (cCRE) or structural-like (CTCF-only) regions in melanoma states, siNTC and siMS. Each dot represents a biological replicate. **c.** Aggregate region analysis at melanocytic, intermediate and mesenchymal CTCF peaks in INT and MES cells and their comparison against MEL, measured as Log_2_ Fold Change of observed/expected HiC contacts. Span and center areas used for statistical comparisons are marked by dotted black squares. **d.** Center and span areas quantification and statistical analysis by one-way ANOVA in MEL, siNTC, siMS, INT and MES cells at melanocytic, intermediate and mesenchymal CTCF peaks. Each dot represents a biological replicate. **e.** Peak annotation of CTCF regions specific of MEL and MES states. **f.** MITF/SOX10^46^ and FOLS2/TEAD4^136^ occupancy at CTCF MES-specific regions. **g.** Immunoblot of CTCF, FOSL1, FOSL2, JUN and pan-TEAD co-immunoprecipitations using MM099 nuclear insoluble extracts. 1% of input and IgG were used as internal controls. Relative molecular weight are shown on the right. **h.** Volcano plots representing the differential binding of transcription factors predicted by TOBIAS footprint analysis. Colored spots are the significantly enriched motifs in the colored matched conditions. Some of the motifs are shown. **i.** siCTCF RNAseq volcano plot of DEGs found downregulated or upregulated in MM099. In red are the genes statistically significant (adjusted p-value <0.05). **j.** Gene Set Enrichment Analysis plot of Mesenchymal and Melanocytic Widmer signatures in MM099 siCTCF. Adjusted p-values and normalized enrichment scores (NES) are shown.

Altogether, our data suggest that CTCF partially relocates from TAD boundaries to enhancer– promoter regions in MES cells, where —together with MES TFs AP1/TEADs— it may directly regulate a subset of MES relevant GRNs. Therefore, melanoma state transitions can involve changes in CTCF deposition and activity.

### State-specific chromatin hubs are highly active in mesenchymal cells, functionally involved in invasion and associated with patients’ survival

We demonstrated that melanoma cells states have profound differences in nuclear architecture irrespective of the mutational driver and burden (i.e., CNVs, SVs). These differences are associated with changes in both the transcriptional output and physical properties (i.e., stiffness) tuning metastatic potential. We hypothesized that MES cell increased chromatin flexibility and inter-domain contacts may favor an increase in cis-CREs (cCREs) connectivity and the formation of new chromatin hubs. To examine global connectivity, valid paired interactions were divided in cis-short (distance < 20 Kb), cis-long (distance > 20 Kb), and trans (inter-chromosomal contacts). To account for DNA mutations/arrangements, we ran EagleC^109^ and HiNT^78^ to detect structural variants (SVs) and copy number variations (CNVs) respectively; valid DNA pairs spanning SVs and CNVs breakpoints were excluded. No significant variations in the number of CNVs and SVs were detected among states and upon siMS (**Supp Fig. 7a-b**).

**Figure 7:**
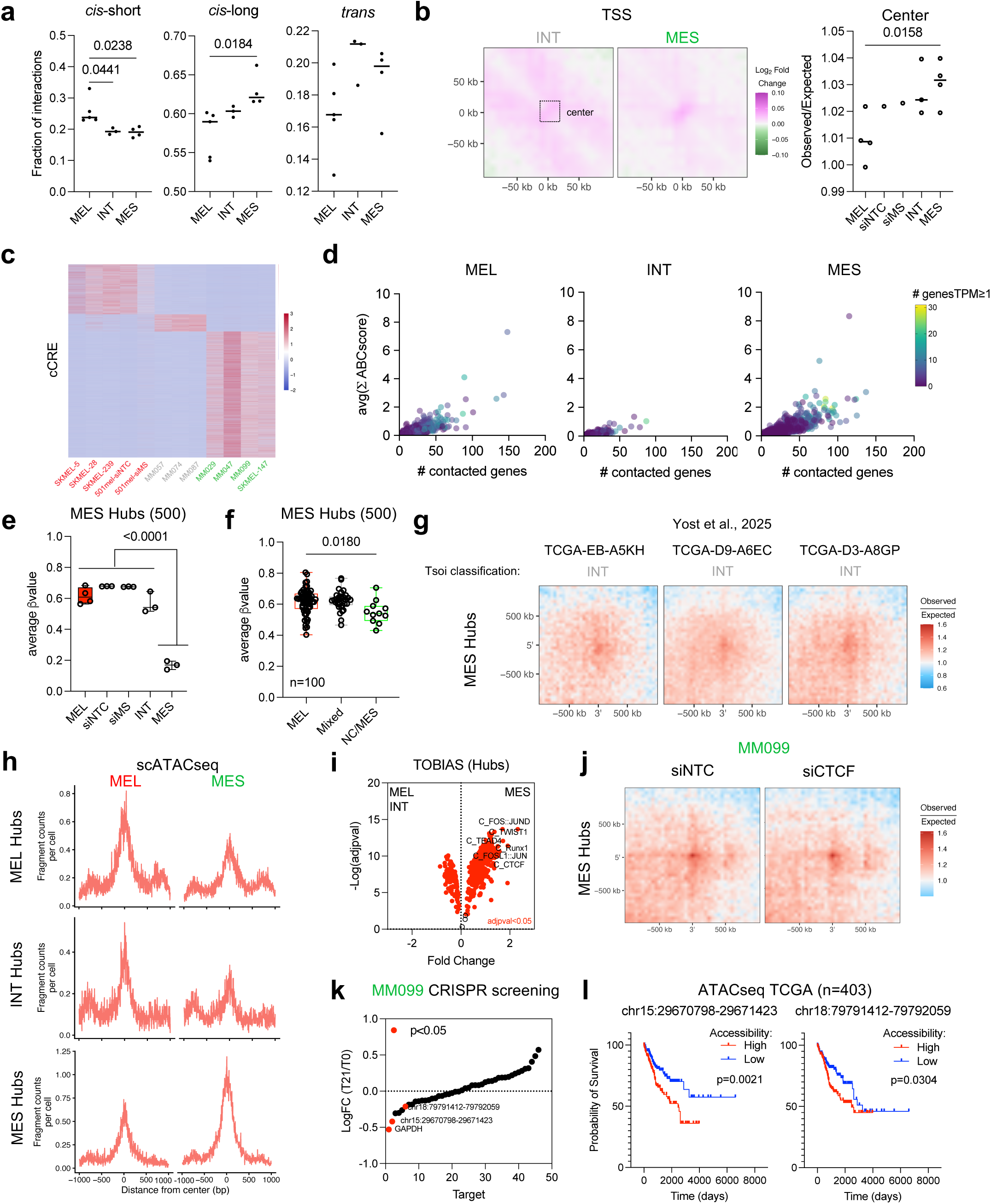
Mesenchymal cells nuclear organization shape chromatin hubs important for migration and cell identity. **a.** Quantification of cis-short (<20Kb), cis-long (>20Kb) and trans HiC contacts by column dot plot in MEL, INT and MES cell lines. One-way ANOVA test was performed, significant p-values are shown. Each dot represents a biological replicate. **b.** (Left) HiC pileup region analysis of genome-wide TSS in INT and MES relative to MEL cells. (Right) Corresponding quantification at the center of the heatmap marked by the dotted black square area. Each dot is a biological replicate statistically evaluate by one-way ANOVA. **c.** Heatmap of the cCRE differentially active in melanoma states. Color scale measure ABCscore z-score normalized per row. **d.** Dot plot distribution of differentially active cCRE in MEL (n=11,581), INT (n=3,987) and MES (n=29,189) cells based on cCRE average sum of ABCscore relative to the number of contacted genes. Color bar measure the number of genes with a TPM expression ≥1 contacted by each cCRE. **e.** Methylation quantification of 500 MES hubs in melanoma cell lines. Methylation levels were averaged from all 500 hubs and one-way ANOVA test was conducted. **f.** Bar dot plot showing the average methylation of 500 MES hubs in primary melanoma samples (n=100) classified as MEL (melanocytic), mixed and NC/MES (neural-crest, mesenchymal). Each dot represent an independent biological sample, one-way ANOVA p-value comparison MEL vs NC/MES is shown. **g.** H3K27ac Hi-ChIP contact aggregate analysis at MES hubs (n=500) in three melanoma patients from the TCGA consortium generated by Yost et al.^98^ and classified in INT based on Tsoi et al.^51^. **h.** Pseudobulk ATACseq enrichment at MEL, INT and MES hubs regions in MEL and MES cells identified by loHiC in 12-273BM derived xenograft. **i.** Volcano plot of the differential TF motifs activity in MES vs MEL/INT hubs according to TOBIAS. Some relevant motifs are shown, in red are the statistically significant motifs found. **j.** HiC aggregate contact analysis at MES hubs (n=500) in MM099 siNTC and siCTCF. **k.** Dot plot representing the CRISPR interfering MES hubs functional ranking in MM099 based on their fold change enrichment from time zero (T0) to 21 days (T21) upon silencing. In red are marked significantly dropped targets. GAPDH serves as internal positive control. **l.** Kaplan-Meier showing the overall survival of TCGA patients (n=404) based on their levels of ATACseq accessibility at two MES hubs found significantly depleted by a CRISPR screen. 50% quartile was used to segregate high (n=202) vs low (n=202) accessibility. Mantel-Cox test p-values are shown.

In line with our hypothesis, MES cells have a significant increase in cis-long range interactions compared to INT and MEL cells, and a reduction in *cis*-short range contacts (**Fig. 7a**). However, siMS treatment does not result in significant variations in global contacts (**Supp. Fig. 7c**). Moreover, HiC pileup map analysis at global TSS shows an augmented connectivity at promoters in MES cells (**Fig. 7b**), an indication of enhanced regulatory activity, which is not mimicked by siMS (**Supp. Fig. 7d**). Next, to map the global cCRE connectivity in melanoma states at the granular level, we applied the previously described ABC-model^82^ to our panel of cell lines, which identified 215,858 cCREs after excluding self-promoter contacts and retaining cCREs with an ABCscore ≥ 0.02 (**Supplemental Table 5**).

To identify state-specific cCREs, we performed DESeq2 analysis of cCREs based on the ABCscore among the states to identify 11,581 MEL-specific cCRE, 3,987 INT-specific cCRE and 29,189 MES-specific cCRE (**Fig. 7c, Supp. Fig. 7e**). Within the state-specific cCRE, we identified chromatin hubs, defined as cCREs able to connect multiple genes concurrently. We ranked cCREs based on the number of genes contacted, the sum of ABCscore provided and the number of contacted genes with a TPM expression ≥ of 1, identifying hubs specific of each state (**Fig. 7d, Supp. Fig. 7f**, **Supplemental Table 6**). In line with our hypothesis, MES cells show the greatest number of hubs compared to INT and MEL. State-specific hub anchors display significantly lower DNA methylation in the corresponding state, suggesting they are epigenetically active (**Fig. 7e**). Reduced DNA methylation was also observed at MES hubs in MES-like specimens of both primary and metastatic melanoma (**Fig. 7f, Supp Fig. 7g**) and this reduction can be detected in patients’ samples^98^ (**Fig. 7g, Supp. Fig. 7h-i**), suggesting that these hubs are active in patients’ tumors and from early tumor stages. To further support the in vivo conservation values of the identified hubs, 12-273BM derived xenograft cells processed with loHiC (**Fig. 4a**) show ATAC accessibility enrichment at hubs of the corresponding identified states (**Fig. 7h**). Next, DNA footprinting of hub anchors revealed a significant enrichment in AP1/TEAD transcription factors as well as CTCF (**Fig. 7i, Supp. Fig 7j, Supplemental Table 7**). Accordingly, MES hub connectivity and activity are significantly affected by CTCF silencing (**Fig. 7j, Supp. Fig. 7k**), further supporting CTCF-AP1/TEAD complexes as key regulators of MES specific GRNs.

Finally, we assessed the functional role of MES chromatin hubs by a drop-out screen using a custom pooled CRISPR interference library against 43 top MES hubs. To test which chromatin hubs are important for MES identity and migratory capacity, MES cells were transduced with a doxycycline-inducible dCas9-KRAB-MeCP2 and the CRISPRi library and seeded for migration on 3.0μm pore transwell inserts. Post-migration cells were collected and re-seeded on a transwell assay multiple times. Twenty-one days after consecutive migration cycles, DNA sequencing revealed two hubs being significantly depleted by all 5 independent sgRNAs/hub (p-value<0.05) compared to time zero (t0) (**Fig. 7k**). Strikingly, ATAC-seq accessibility of these two hubs is significantly associated with shorter patient survival across various cancer types^110^ (**Fig. 7l**).

Together, our results precisely map the most active cCREs in the main melanoma phenotypic states. Particularly in MES cells, it revealed the bivalent function of chromatin architecture in which CTCF gets recruited at hubs and cCREs of genes involved in MES identity at the expense of domains’ boundaries, simultaneously promoting a more flexible nuclear architecture which favors cell migration and intravasation.

## Discussion

State transitions and phenotypic plasticity have been linked to cancer metastasis and therapeutic resistance. In particular, the acquisition of a de-differentiated, mesenchymal identity is a common feature across tumors of various origins, such as carcinomas^111, 112^, glioblastoma^113^ or melanoma^114^. To date, most studies have attributed associated transcriptional programs with confer cancer cells with a selective advantage under multiple stressors as they invade, colonize distal tissues, or survive treatment. While previous studies have been limited to a two-dimensional analysis of state-specific epigenomes, we are the first to dissect the contribution of higher order chromatin structures and nuclear architecture to mesenchymal (MES) identity, using melanoma as a paradigmatic model of metastasis. Contrary to differentiated MEL state, we demonstrate that the MES state encompasses a loose chromatin architecture and nuclear conformation which supports not just a distinct transcriptional and epigenetic program but mechanical properties such as nuclear deformability as an adaptive response to migratory constriction and metastasis. MES nuclear topology results in part from a relocation of CTCF from TADs boundaries to MES-specific chromatin hubs and cCREs, and enhanced cis-long interactions (larger TADs, more inter-TADs contacts) and reduced H3K9me3 constitutive heterochromatin (small, immature CPDs) relative to differentiated, MEL cells. MES cells represent an intriguing, de-differentiated cancer cell state conserved across multiple cancer types^115^. Our data suggests that multiple tumor types from various cellular origin may similarly converge from a differentiated state, with a ‘rigid’ chromatin architecture locked by lineage-specific GRNs, to a pan-cancer undifferentiated (MEL-like) state, broadly characterized by loose chromatin, nuclear elasticity and enhanced deformation upon constriction, as we demonstrate in melanoma.

Supporting this model, silencing of the master melanocytic TFs MITF and SOX10 GRNs can ‘push’ melanoma cells toward an intermediate (INT-like) state as previously reported^46^, but this is insufficient in allowing cells to adopt MES nuclear architecture and mechanical features, which suggests the requirement of additional stimuli. We showed that upon physical constriction such as the one experienced during migration, melanocytic cells shift to a MES-like nuclear molecular conformation, characterized by reduced CTCF occupancy and insulation at TADs’ boundaries, lower heterochromatin and stiffness, and decreased MITF and SOX10 levels, which confers them with enhanced metastatic potential. This bidirectional crosstalk between 3D chromatin and nuclear compression links for the first time epigenetic features, gene expression programs and mechanical properties to cancer phenotypic plasticity and metastasis.

Traditionally, cancer cell phenotypes have been simply defined by differential transcriptional states^116^, where lack of genetic drivers indicates that, at least in part, they are epigenetically driven^117, 118^. However, the role of epigenetic regulatory mechanisms in defining these transcriptional states remains poorly understood, partly due to the limited application of in vivo epigenetic technologies at single-cell resolution (mostly ATACseq)^119^. Although some state-specific gene’ functions clearly map to a corresponding state (e.g., ATF4 in stress response, E2Fs in mitosis, MMPs in invasion, PD-L1 in immune evasion)^43^, many others are harder to interpret because their expression overlaps across states^116^. Instead, these overlapping genes may reflect indirect downstream adaptations to upstream epigenetic changes, such as alterations in chromatin architecture, rather than representing state-specific functional programs. In support of this, we generated the first atlas of melanoma cell state regulatory element (n=215,858) activity and their connections with other genes, which can inform future cancer epigenetic-related work. Here, melanoma states exhibit profound, global differences in 3D chromatin organization that are not explained solely by differential gene expression^120^, but also by distinct nuclear organization and physical properties *in vitro* and *in vivo*. These results contribute to the ongoing debate on the role of chromatin architecture in global versus gene-specific transcriptional regulation^121^. Our findings support a model of converging, transcription-dependent and independent roles of chromatin architecture, exemplified by the repurposing of CTCF as both a structural and transcription factor, and the emergence of chromatin hubs as regulatory network organizers. Untangling CTCF gene regulatory function is demanding due to its polyhedric role as activator, repressor and genome organization protein^122^, as shown in mouse embryonic stem cells where it acts as activator of a subset of genes together with its paralog CTCFL^123^. In cancer cells, CTCF can shift from a structural to a gene-regulatory role, as seen in MES cells where it binds less to insulators and more to CREs, promoting *cis*-long interactions, inter-TAD connectivity and chromatin hubs formation, regulatory headquarters required for the expression of cancer cell identity genes^23, 24, 57^. In agreement, a brief depletion of CTCF affects MES hub connectivity and their target genes, and our CRISPRi screening revealed a previously unreported role of hubs in controlling MES cells’ biology. The two identified hubs control 72 genes involved in cell proliferation (i.e., NCAPGP2, CTDP1, PARD6G)^124–126^, migration (i.e., NFATC1, TJP1)^127, 128^ and survival (UBE3A)^129^. Yet, it is unclear if chromatin hubs solely control gene expression or also nuclear flexibility. In fact, although physical constriction is sufficient to reduce CTCF at MEL boundaries, it is insufficient to recruit it at MES CREs and hubs, which may require TF partners^120^ such as AP1/TEADs and CTCFL, all of which have been shown to promote a MES state in melanoma^47, 130–132^.

In addition, we demonstrate that MES cells display a reduced maturation and size of physical CPDs, which directly correlates with mechanical stability. For example, stable mature domains have been shown to lower the probability of blebbing during cell migration, with predominantly ‘domain-less’ chromatin forming the blebs.^34^ In addition, mechanical stressors and forces acting on cancer cells are expected to affect CPDs and their encoded transcriptional memory through a selective process favoring immature and smaller CPDs (low H3K9me3) rather than mature (high H3K9me3). Due to their greater steric hindrance, increased crowding upon constriction can convert mature CPDs into immature ones, destabilizing H3K9me3, CBX5 (HP1a)^13, 14, 133^ and CTCF insulation which is required to contain heterochromatin nucleation^12^. Cell differentiation requires H3K9me3 deposition to silence lineage unrelated genes expression^134^ within mature CPDs to allow transcriptional memory^16, 101^. Therefore, MES cells may represent a phenotypic state in which mechanical confinement, coupled with the loss of lineage-determining GRNs (e.g., MITF and SOX10 in melanoma), leads to erosion of the transcriptional memory that encodes tumor lineage identity (i.e., the melanocytic-like state), as reflected by CPD immaturity, increased long-range cis contacts, TAD expansion, and reduced CTCF-mediated insulation. Based on this model, cells in the MES state may lose plasticity by efficiently propagating a stabilized epigenetic and 3D chromatin configuration through cell division, resulting in maintenance of MES identity during tumor progression. This is consistent with observations in glioblastoma, where the MES phenotype is highly heritable across cell generations^7^. Similarly, while MEL to MES dedifferentiation has been extensively reported^51, 135^, there is little to no evidence for the opposite process (i.e., from a de-differentiated to a differentiated state).

Finally, our study provides the first evidence of a direct link between 3D chromatin and nuclear biophysical properties and cancer cell states in actual tumor samples. By combining state-of-the-art bulk and single cell high-resolution microscopy (PWS, sSMLM), epigenomics and molecular and cell biology techniques, we demonstrate their potential for clinical translation. For instance, we integrate PWS and multiplex IF (Opal) staining-which are relatively inexpensive and rapid techniques-on a curated TMA containing. Our future work will integrate 3D genome and nuclear-property data across larger cohorts to train AI models on H&E images, enabling machine-learning–based identification of cancer cell states and linking intra-tumor heterogeneity to patient outcomes. Our findings identify nuclear chromatin organization as an essential driver of cancer cell plasticity, representing a novel intersection between mechanical properties, epigenetics and gene expression in tumor progression.

## Acknowledgments

This work was supported by U54CA263001 (E.H., I.O and A.W.L.) and R01CA274100 (E.H.), from NCI/NIH. P.B. was awarded of a National Cancer Center (NCC), Melanoma Research Alliance (MRA) and Melanoma Research Foundation (MRF) fellowships. We thank the Genomics Technology Center (GTC), particularly Mr. Paul Zappile, for supervising all the sequencing; the Preclinical Imaging Core (Mr. Orlando Aristizabal), the Experimental Pathology Core (Dr. Cynthia Loomis and Mr. Gyles Ward for Opal panel optimization) and the Flow Cytometry core, all of which are supported by Cancer Center Support Grant (CCSG) from NCI/NIH to the NYULH Perlmutter Cancer Center (P30CA016087; PI: Alec Kimmelman). V.B. was supported by U54 (CA268084) and L.A. by a K23 (1K23DK144661) NIH/NCI awards. This research used resources of the Center for Physical Genomics and Engineering at Northwestern University, and philanthropic support from K. Hudson and R. Goldman, S. Brice and J. Esteve, M. E. Holliday and I. Schneider, the Christina Carinato Charitable Foundation, and D. Sachs. J.S. was supported by R35GM122525, and J.S. and I.A. by P01CA229086 NIH/NCI grants. T.M. was granted a Cancer Research Institute Dr. Keith Landesman Memorial Postdoctoral Fellowship (CRI14497).

## Author contributions

P.B. and E.H. conceived and design the work. P.B. performed and analyzed most of the omics experiments (HiC, ChIPseq, ATACseq, loHiC, CUT&Tag, scRNAseq, CRISPR screen, Methylation, Polychrom model and public datasets mining). V.B, C.D. and L.A. designed and conceived the microscopy and biophysical experiments, and C.D. and L.A. performed the experiments. K.I.M and N.A. performed PWS and sSMLM imaging, analysis and visualization. J.S. and C.Do provided helpful insights for the HiC analysis. A.F.Y. and P.B. performed the chromatin hubs CRISPR screen and analysis, and *in vivo* mouse experiments. I.A. and S.L. shared the loHiC protocol and S.L. provided analysis support. I.O. and M.I. provided patient data and material under IRB protocol #H10362 and designed the TMA. A.S.M. provide a pathology evaluation of the TMA. T.M. analyzed the TMA staining. S.N.E., D.C. and N.E.T. provided the cell line methylation data. A.R., C.K. and R.S. provided the methylation data of a cohort of primary melanoma of the international consortium InterMEL. S.N.E., D.C., N.E.T., K.D., A.R., C.K., R.S. and E.H. are part of InterMEL. P.B., E.H., C.D., L.A. and V.B. interpret the results. P.B. and E.H. wrote the manuscript with important inputs from C.D., L.A. and V.B. All authors approved and revised the final manuscript.

---

**Supplemental Figure 1:**

**a.** Snapshot captured by confocal microscopy showing the staining of different markers using Opal in cells classified as MEL-like and MES-like found in two melanoma TMAs.

**b.** sSMLM images of H3K9me3 and H3K27ac nuclear distribution in SKMEL-5 (MEL), MM057 (INT) and MM099 (MES).

**c.** Average H3K27ac distribution genome-wide in MEL, INT, MES. Each profile represents the average of at least three independent cell lines.

**d.** H3K9me3 staining in 501mel siNTC and siMS captured by sSMLM.

**e.** H3K9me3 and H3K27ac ChIP-seq genome-wide distribution in 501mel siNTC and siMS.

**Supplemental Figure 2:**

**a.** Normalized genomic occupancy (RPKM) of H3K9me3, H3K27ac, ATAC and RNA at active (A) and inactive (B) compartments in MEL, INT, MES, siNTC and siMS cells. Standard deviations bars are based on individual compartment BIN of 100kb spanning the whole genome. Unpaired t test was performed comparing A vs B per each marker (H3K9me3, H3K27ac, ATAC and RNA).

**b.** Grouped column bar dot plot of the percentage of A and B compartments in *cis* and *trans* identified in melanoma states (top) and siNTC vs siMS (bottom). Two-way ANOVA test was performed to find significant differences. Each dot represents a biological replicate.

**c.** HiC *trans* compartment saddle plots of melanoma states (top) and siNTC vs siMS (bottom). Values of 20% quantile areas are shown.

**d.** Quantification of trans compartment AA, AB:BA and BB interactions shown in panel c. Each dot is a biological replicate. Unpaired t-test p-value of BB interactions in siNTC and siMS is shown.

**e.** Column dot plot showing the RPKM enrichment of H3K9me3, H3K27ac, ATAC (left) and RNA (right) at compartments switching from A to B or B to A found in compartment switch analysis between states, siNTC and siMS. (Left) Each dot represent a single compartment of 100kb BIN. (Right) Each dot represent the average of normalized RNA reads, standard deviations bars are generated by compartments replicates. Unpaired t-test comparison was used. Significant p-values are displayed.

**Supplemental Figure 3:**

**a.** Insulation score metaprofiles at MEL, INT and MES TADs boundaries across all melanoma cell lines.

**b.** CTCF occupancy at MEL, INT and MES TADs boundaries in all cell lines. For a-b panels, red, grey and green colors mark melanocytic, intermediate and mesenchymal cells respectively.

**c.** Insulation (top) and CTCF (bottom) profiles at siNTC, siMS, MEL, INT and MES TADs boundaries.

**d.** HiC aggregate contact analysis of siMS cells at both siNTC and siMS TADs relative to siNTC cells.

**e.** Relative intra-TAD (0) and inter-TAD (+1,+2,+3) HiC contacts in siMS relative to siNTC.

**f.** Mean CTCF distribution in siNTC and siMS cells at MEL, INT, MES, siNTC and siMS loops.

**g.** HiC aggregate peak analysis at MEL, INT and MES loops anchors in siMS, INT and MES cells relative to MEL/siNTC cells.

**h.** Percentage of TADs (left) and loops (right) that are found common (shared) or specific of each state, siNTC and siMS. **i**. Volcano plot showing the DEGs found upon siMS in 501mel. MITF and SOX10 are shown. In red are DEGs with a significant adjusted p-value.

**Supplemental Figure 4:**

**a.** Signac ATAC accessibility at *MITF* (top) and *EGFR* (bottom) loci in 12-273BM melanoma states identified by loHiC.

**b.** HiC contact heatmap from chromosome 1 to X in MEL and MES cells profiled by loHiC. Color scale represents the log1p of contacts.

**c.** Copy number variations (CNVs) analysis plot predicted by HiC in MEL and MES cells profiled by loHiC. Pearson correlation coefficient and p-value comparing CNVs in MEL vs MES are shown.

**d.** Saddle plot of *cis* compartments in MEL and MES cells identified by loHiC.

**e.** Bar dot plot of the HiC contacts intra-TAD (o) and inter-TAD (+1,+2,+3) in MES cells relative to MEL cells. Paired t-test was conducted comparing MEL vs MES at each TAD neighbor.

**Supplemental Figure 5:**

**a.** 3D rendering of chromosome 1 modeled as a polymer chain using polychrom and based on the three principal components. Each bead is a 10kb BIN color scaled based on the predicted bead average softness. In grey are beads falling into blacklisted regions. The average chromosome 1 radius of gyration (Rg) per cell line are shown.

**b.** Quantification of Rg changes upon a simulation of constriction of chr1 polymer chain. One-way ANOVA test was performed. Each dot represent a biological replicate.

**c.** Column dot plot quantify the z-score enrichment of melanocytic and mesenchymal Widmer signatures^53^ in 501mel Pre and Post migration. Unpaired t-test comparing Pre and Post for each signature was performed. Each dot is representative of an independent biological replicate.

**d.** *cis* compartments saddle plot of Pre and Post 501mel and MM099 cells. 20% quantiles are shown.

**e.** Scatter plot of compartment switching between Pre and Post cells in 501mel (top) and MM099 (bottom). In black are highlighted significantly switching compartments.

**f.** Melanocytic and mesenchymal Widmer signatures enrichment in MM099 Pre and Post quantified as averaged z-score. Three biological replicates were used. Unpaired statistical t-test was conducted.

**g.** Chromatin diffusion (D_e_) quantification in 501mel and MM099 Pre and Post migration. Unpaired t-test was conducted in Pre vs Post and Pre vs Pre pairwise comparisons. Resulted p-values are shown.

**h.** Representative images captured by sSMLM for H3K9me3 and H3K27ac staining in MM099 Pre and Post migration.

**i.** Proliferation curve of 501mel Pre and Post measured as percentage of well confluence from time 0 till 96hrs. Each dot is the average of three biological replicates. Unpaired t-test was conducted to compare the end point.

**j.** Snapshot of GFP fluorescence representing metastatic foci in livers of NSG injected intracardiac with 501mel Pre (n=7) or Post (n=7).

**k.** (Left) Representative images of SKMEL-28 (MEL) post migratory cells upon 12hrs of DMSO (control) or Chaetocin treatment. (Right) SKMEL-28 post migratory relative quantification to the DMSO. Each dot represent an independent biological replicate. Paired t-test was performed. P-values is shown.

**l.** Immunoblot showing H3K9me3, MITF and SOX10 variation upon 24hrs DMSO or Chaetocin (CHAE) treatments in 501mel and SKMEL-28. H3 and VINCULIN serve as internal loading controls. Relative molecular weights are shown on the right.

**Supplemental Figure 6:**

**a.** (Left) Immunoblot for CTCF, MITF and EGFR across multiple melanoma cell lines used in this study. Vinculin serve as loading controls. Relative molecular weights are shown on the left. MEL, INT and MES cells are labeled in red, grey and green respectively. (Right) CTCF protein quantification based on the immunoblot shown on the right. One-way ANOVA test was used to assess statistical significance between MEL, INT and MES cells. Each dot represent a protein band of a biological replicate.

**b.** Aggregate region analysis of siMS HiC contacts relative to siNTC at CTCF bound regions of MEL, INT and MES cells. **c**. CTCF state-specific regions identified with Diffbind and profiled for CTCF in all cell lines as averaged profile (top) and heatmap (bottom).

**c.** Pie chart representing the percentage of CTCF MES-specific regions (n=1,274) annotated as CTCF-only (structural), intragenicEnh (intragenic enhancer), intergenicEnh (intergenic enhancer) or promoter.

**d.** ABCscore quantified with a column dot plot. Each dot represent the average ABCscore value of cCREs (n=1,479) identified with the ABC model found to overlap with CTCF MES-specific regions. One-way ANOVA test was conducted, a summary p-value of the comparison MES vs all is shown.

**e.** GSEA enrichment plot of the 5,081 genes found to be contacted by CTCF in MES cells and scored in Tsoi et al.^51^ short-term culture cells classified as MEL (n=17) and MES (n=10). NES and adjusted p-values are shown.

**f.** 5,081 genes signature score (ABC_5k) in melanocytic and mesenchymal cells found in melanoma patients by Pozniak et al.^85^ Unpaired t-test p-value is displayed.

**g.** Immunoblot for CTCF and ACTB (loading control) in whole cell protein extracts of MM099 siNTC and siCTCF cells. Relative molecular weights are shown on the right.

**h.** Venn diagram of the overlap between significantly DEGs siCTCF and ABC-5k. Hypergeometric test was conducted, representation factor and p-value are shown.

**i.** Violin plot of the downregulated (Down) and upregulated (Up) genes upon siCTCF in melanocytic and mesenchymal melanoma cell states of patients’ samples^85^. Unpaired t-test p-values are shown.

**Supplemental Figure 7:**

**a.** Quantification of the number of CNVs as a grouped column plot across MEL, siNTC, siMS, INT and MES samples. Two-way ANOVA test was conducted.

**b.** Column dot plot of the number of structural variants (SVs) found in the melanoma cell lines used in this study. One-way ANOVA test was performed.

**c.** Fraction of *cis*-short, *cis*-long and *trans* HiC contacts in siNTC and siMS cells. Paired t-test was used.

**d.** HiC aggregate region analysis at all TSS in siMS relative to siNTC.

**e.** PCA plot of the most variable cCREs identified with DESeq2.

**f.** ABC quantification of the top 500 state-specific cCRE Hubs across all cell lines. Bar plots show the average of all hubs. Standard error of the means bar are shown. Red, grey and green mark melanocytic, intermediate and mesenchymal cell lines respectively.

**g.** Bar dot plot quantify the average methylation of 500 MES hubs in TCGA melanoma patients (n=470) classified in MEL, INT, NC (neural crest) and MES according to Tsoi^51^. One-way ANOVA p-value summarizing NC and MES vs INT and MEL comparisons is shown.

**h.** HiC aggregate pile up heatmap of contacts at MES hubs in FFPE patients’ specimen classified in MEL (n=3) and MES (n=4).

**i.** Quantification of HiC contacts at the foreground of MEL and MES hubs in MEL and MES melanoma patients (n=7). Unpaired t-test p-values are shown.

**j.** CTCF average genomic occupancy at MES hubs (n=500) in individual mesenchymal cell lines.

**k.** Volcano plot of the differential MES cCREs activity in siCTCF MM099 cells predicted by RNAseq. In blue are marked the significantly dysregulated cCREs, in red the hubs. Y-axis dotted line show the significant p-value threshold. On top are marked the number of significantly active MES hubs (42 down, 136 up). Multiple t-test with Benjamini/Hochberg correction was used.

For panels a,b,c,e,g and j, each dot is representative of a biological replicate.

## Supplementary information and material

**Supplemental Table 1:** Clinical and molecular information of the tumors’ cores of the tissue microarray.

**Supplemental Table 2:** Differential expression analysis outputs in melanocytic, intermediate, mesenchymal and siMS. In yellow are marked the significantly positive associated gene to the corresponding condition.

**Supplemental Table 3:** Clinical and pathological information of the FFPE blocks processed for HiC.

**Supplemental Table 4:** List of genes contacted by CTCF in MES cells predicted by the ABC model.

**Supplemental Table 5:** List of all cCREs identified by the ABC model, their target genes and activity in melanoma cell states.

**Supplemental Table 6:** Ranked list of cCREs found significantly associated to MEL, INT and MES cells.

**Supplemental Table 7:** Transcription factor motif list analyzed for enrichment at hubs comparing MES vs MEL and INT cells.

**Supplemental reagents:** Complete list of reagents used in this work.

