## Supplemental Figures for "Chromatin architecture and physical constriction cooperate in phenotype switching and cancer cell dissemination"

### Supplemental Figure 1

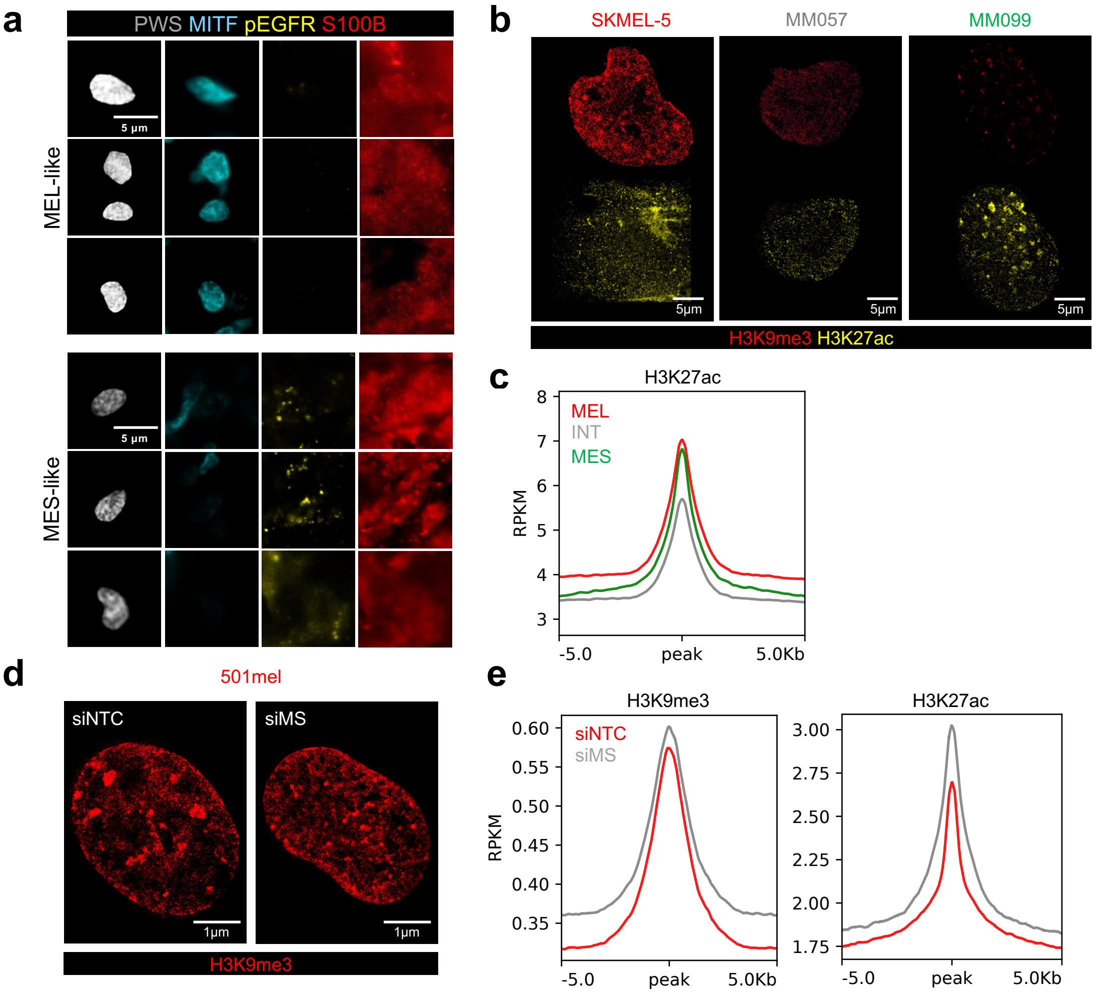

### Supplemental Figure 2

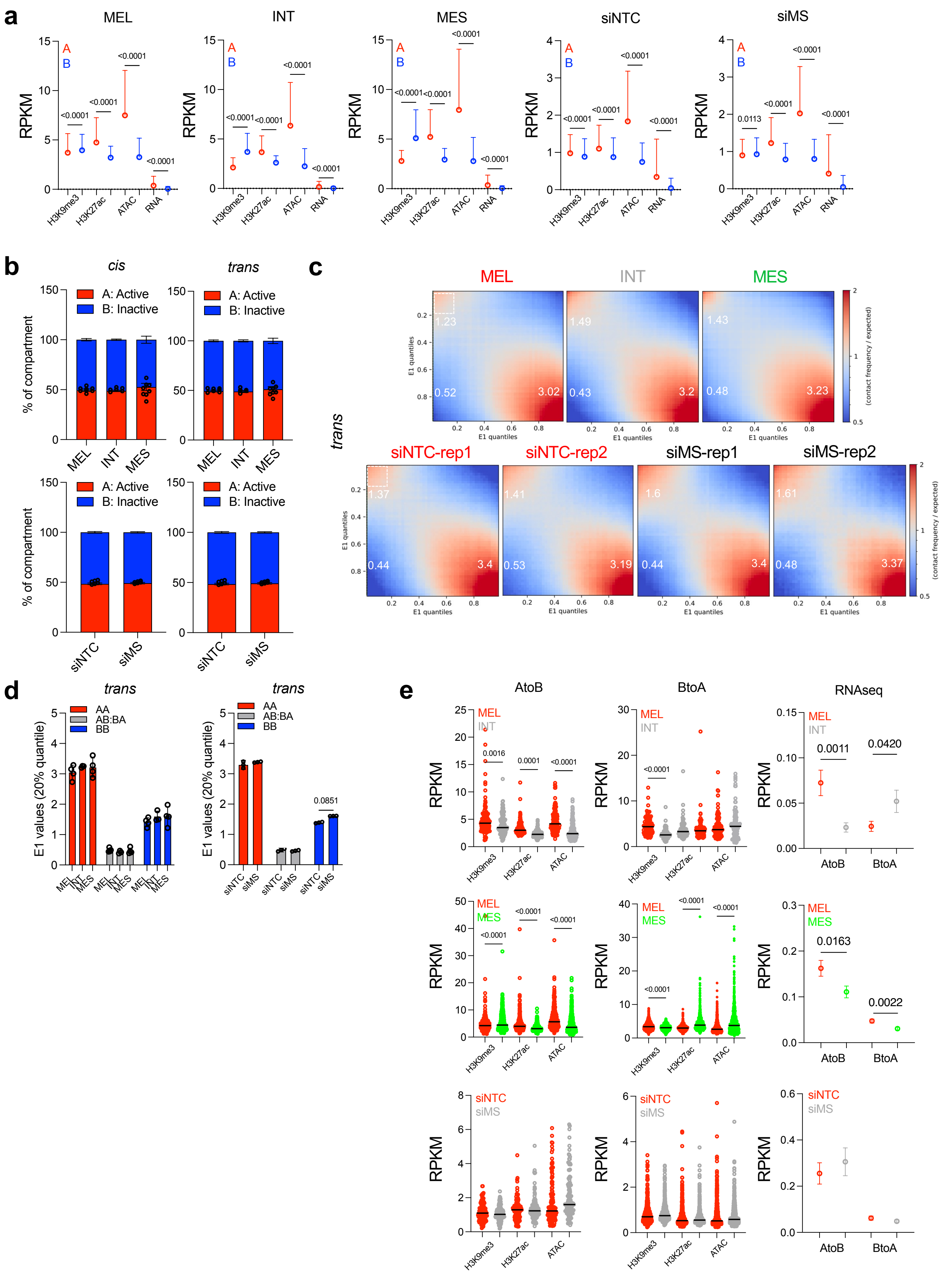

### Supplemental Figure 3

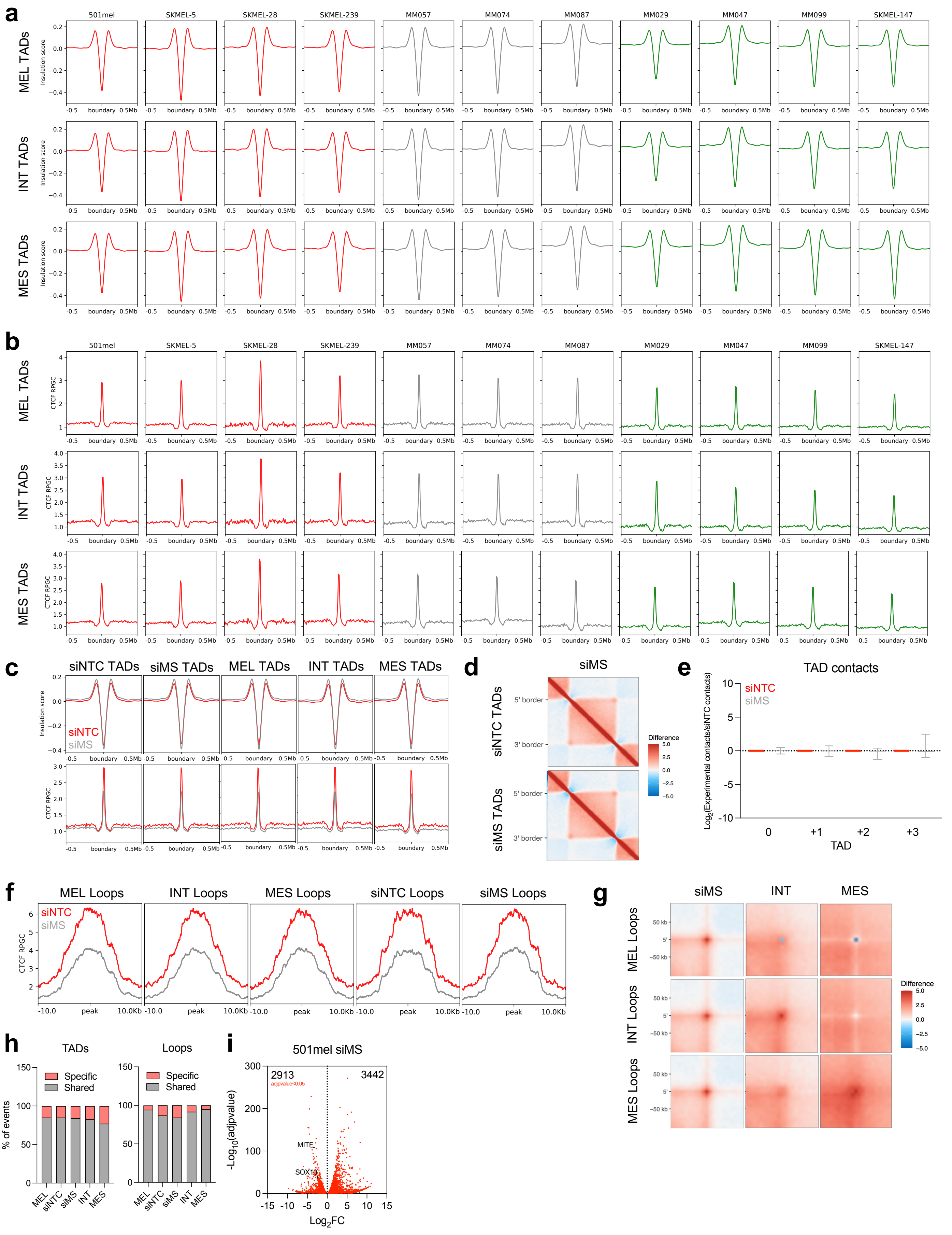

### Supplemental Figure 4

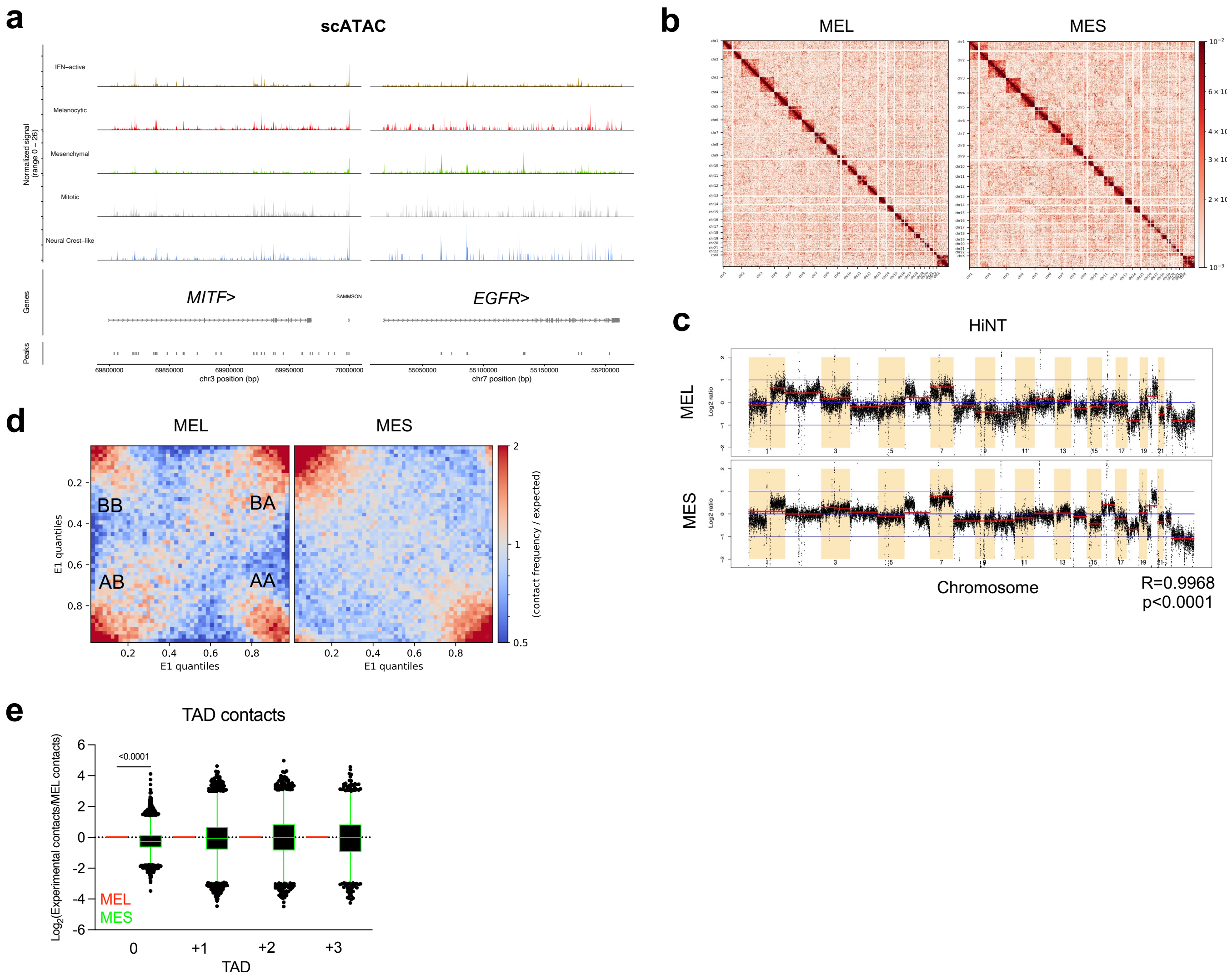

### Supplemental Figure 5

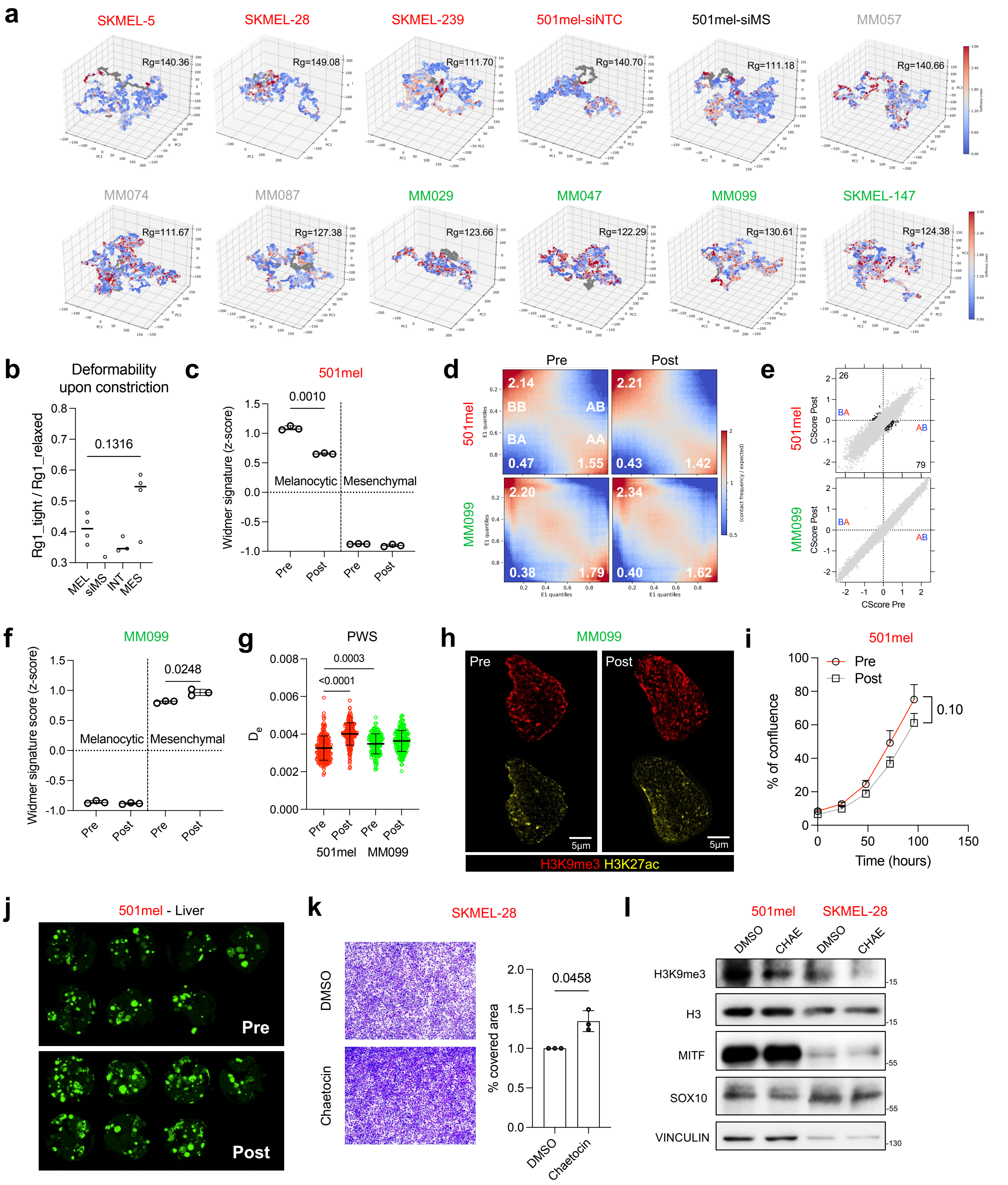

### Supplemental Figure 6

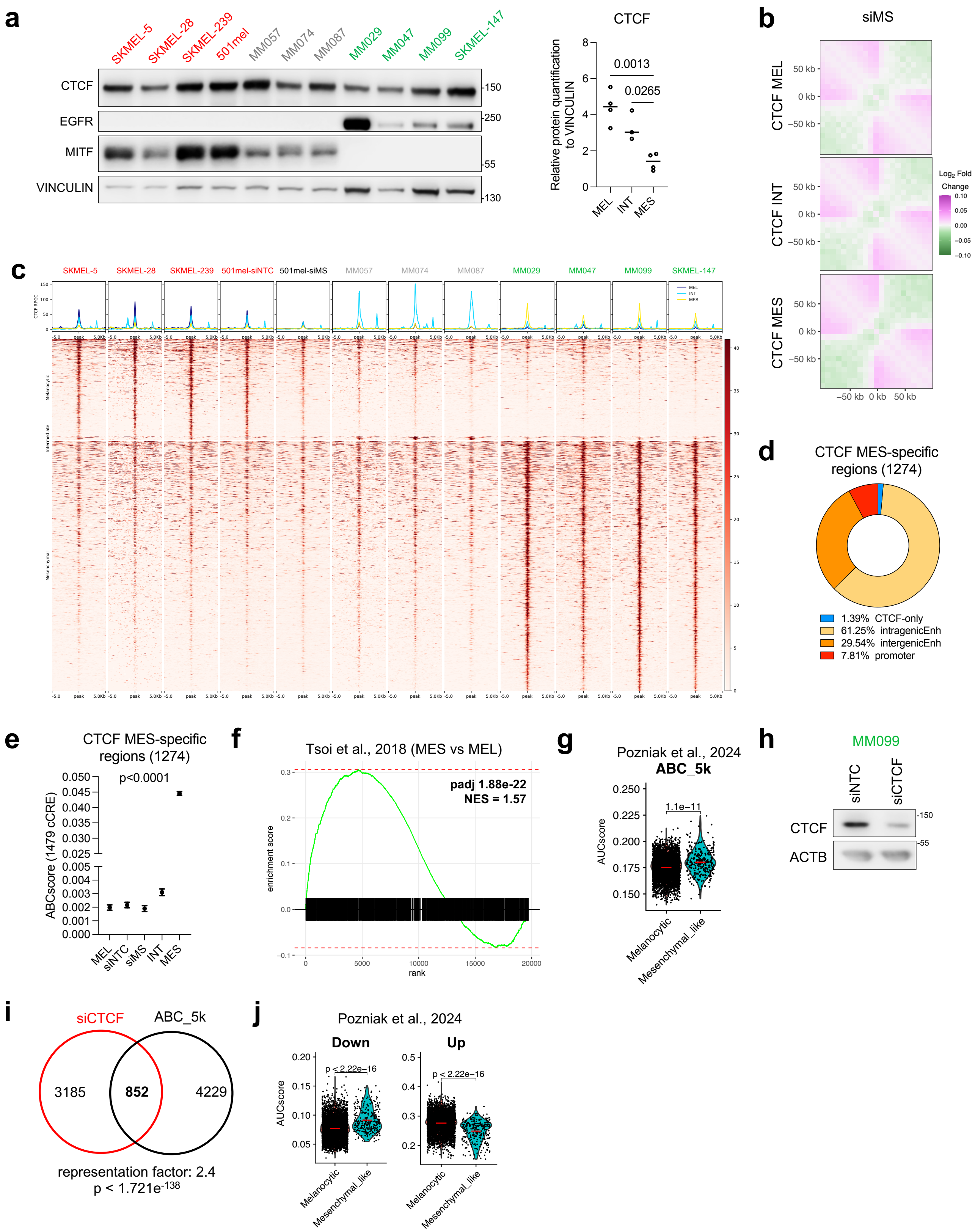

### Supplemental Figure 7

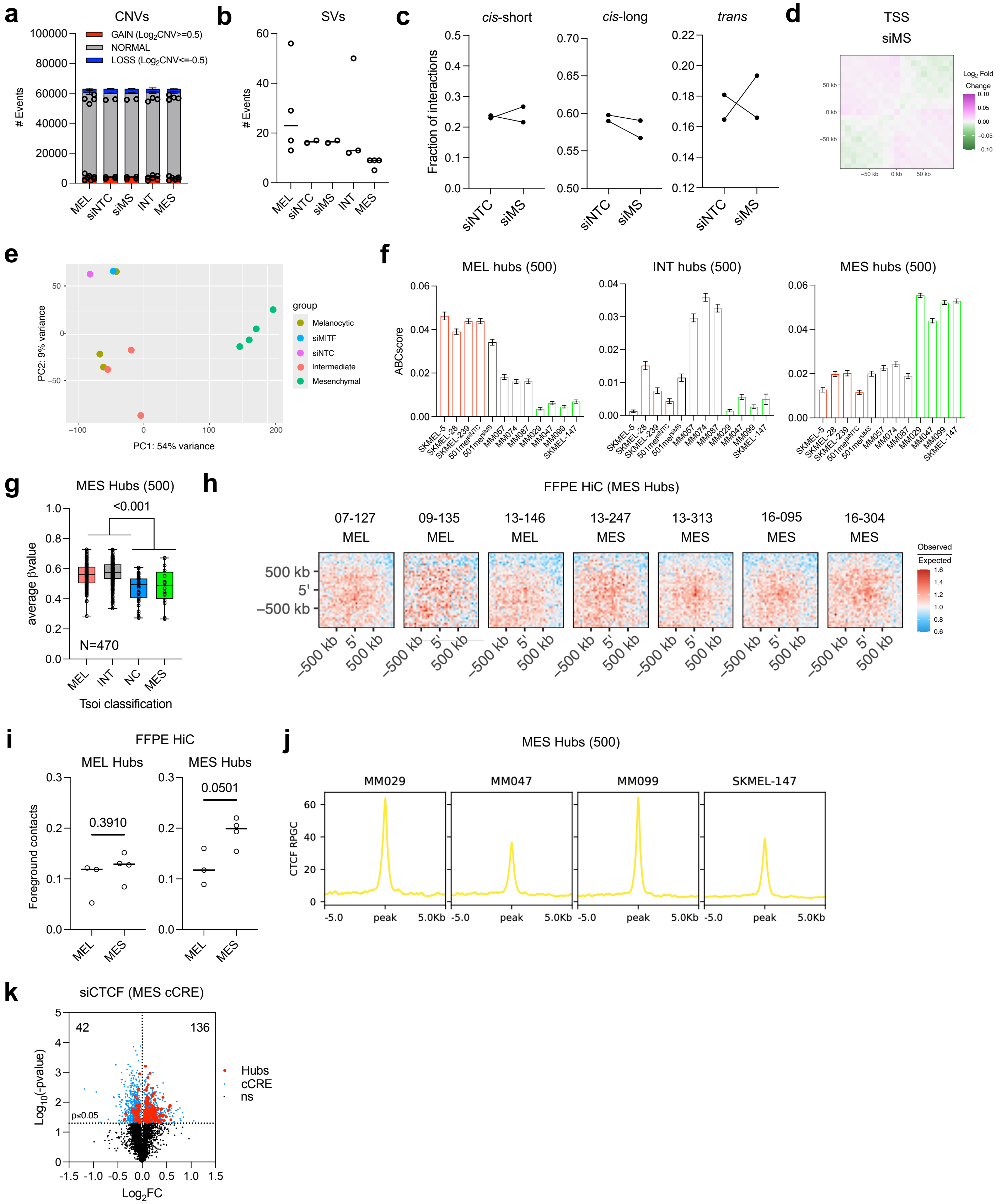
